# Autophagy counters inflammation-driven glycolytic impairment in aging hematopoietic stem cells

**DOI:** 10.1101/2023.08.17.553736

**Authors:** Paul V. Dellorusso, Melissa A. Proven, Fernando J. Calero-Nieto, Xiaonan Wang, Carl A. Mitchell, Felix Hartmann, Meelad Amouzgar, Patricia Favaro, Andrew DeVilbiss, James W. Swann, Theodore T. Ho, Zhiyu Zhao, Sean C. Bendall, Sean Morrison, Berthold Göttgens, Emmanuelle Passegué

## Abstract

Aging of the hematopoietic system promotes various blood, immune and systemic disorders and is largely driven by hematopoietic stem cell (HSC) dysfunction (*1*). Autophagy is central for the benefits associated with activation of longevity signaling programs (*2*), and for HSC function and response to nutrient stress (*3,4*). With age, a subset of HSCs increases autophagy flux and preserves some regenerative capacity, while the rest fail to engage autophagy and become metabolically overactivated and dysfunctional (*4*). However, the signals that promote autophagy in old HSCs and the mechanisms responsible for the increased regenerative potential of autophagy-activated old HSCs remain unknown. Here, we demonstrate that autophagy activation is an adaptive survival response to chronic inflammation in the aging bone marrow (BM) niche (*5*). We find that inflammation impairs glucose metabolism and suppresses glycolysis in aged HSCs through Socs3-mediated impairment of AKT/FoxO-dependent signaling. In this context, we show that inflammation-mediated autophagy engagement preserves functional quiescence by enabling metabolic adaptation to glycolytic impairment. Moreover, we demonstrate that transient autophagy induction via a short-term fasting/refeeding paradigm normalizes glucose uptake and glycolytic flux and significantly improves old HSC regenerative potential. Our results identify inflammation-driven glucose hypometabolism as a key driver of HSC dysfunction with age and establish autophagy as a targetable node to reset old HSC glycolytic and regenerative capacity.

**One-Sentence Summary:** Autophagy compensates for chronic inflammation-induced metabolic deregulation in old HSCs, and its transient modulation can reset old HSC glycolytic and regenerative capacity.

## Main Text

Although different tissues undergo distinct alterations with age, one overarching hallmark of aging is reduced stem cell function in regenerative tissues (*6*). The composition of the blood system evolves with age alongside well-characterized changes in HSC activity, including an age-dependent expansion of the HSC pool with decreased regenerative potential in transplantation assays, a skewed differentiation towards myeloid cell production at the expense of lymphopoiesis and erythropoiesis, and a perturbed state of quiescence characterized by increased stress-response signaling (*6,7*). In addition, aging remodels the BM niche and increases both systemic and microenvironmental inflammation that directly contribute to impaired blood production (*1,5,7,8*). Together, these changes promote the onset of blood cancers, increase susceptibility to deadly infections, and drive chronic inflammatory disorders and systemic tissue degeneration in the elderly (*1*).

HSCs are tightly regulated by their metabolic status, which underlies their sensitivity to metabolic deregulation in aging (*9*). Quiescent HSCs predominantly utilize glycolysis and limit mitochondrial glucose oxidation, whereas activated HSCs increase glucose consumption and link glycolysis with mitochondrial metabolism in order to meet proliferative requirements (*10,11*), as do other proliferating blood cells such as lymphocytes (*12,13*). Genetic loss of glycolysis-promoting regulators or downstream effectors, as well as aberrant oxidative phosphorylation (OXPHOS) activation, prevent maintenance of HSC quiescence and promote stem cell exhaustion (*14,15*). These metabolic states are closely linked to the regulation of nutrient-sensing signaling pathways and the phosphatidylinositol-3-kinase (PI3K)/protein kinase B (AKT)/mammalian target of rapamycin (mTOR)/forkhead transcription factors (FoxO) signaling cascade in HSCs (*16–18*). They are also regulated by fundamental proteostasis mechanisms, particularly macro-autophagy (hereafter called autophagy) that is engaged in HSCs through a FoxO3a pathway to withstand metabolic starvation (*3*) and is used at steady-state to recycle activated mitochondria, manage lysosomal function, and terminate OXPHOS-mediated pro-differentiation signaling (*4,11*). Additionally, asymmetric inheritance of lysosomes and autophagosomes predicts HSC activation state (*19*), and the activity of chaperone-mediated autophagy, a selective form of lysosomal protein degradation that is essential to sustain HSC activation in young mice, decreases over time in old mice (*20*). In contrast, old HSCs remain competent for autophagy induction in stress conditions and, in contrast to many other aging tissues, show increased basal activation of autophagy (*3*). Strikingly, autophagy-activated old HSCs have higher regeneration potential than old HSCs that do not engage autophagy and consequently display overactive oxidative metabolism (*4*). However, which signals promote autophagy engagement in the aged HSC compartment and why autophagy activation increased the regenerative potential of old HSCs remain unknown. Understanding these connections has direct translational implications for restoring old HSC function.

### No identity drift between old HSC subsets

To explore the differences between old HSCs (oHSC) that do or do not engage autophagy, we used ∼ 24-month-old *Gfp-Lc3* autophagy-reporter mice to isolate autophagy-activated (AT^hi^, 33% of cells with the lowest GFP levels) oHSCs based on their characteristic left-shift in GFP-LC3 fluorescence levels that distinguished them from autophagy non-activated (AT^lo^, 33% of cells with the highest GFP levels) oHSCs (**Fig. 1A; fig. S1A,B**). We also used ∼ 2-month-old *Gfp-Lc3* mice to isolate young HSCs (yHSC) without separation based on GFP-LC3 levels. While old HSCs show variations in their degree of autophagy activation, this standardization method and inclusion of yHSC controls allow consistent identification of AT^hi^ and AT^lo^ oHSC subsets between aged mice (*4*). The use of GFP-LC3 signal intensity to monitor autophagy engagement in both young and old HSCs has also been extensively validated in our previous publications (*3,4*). To gain insights into the molecular architecture of AT^hi^ and AT^lo^ oHSCs, we first assessed chromatin accessibility by Assay for Transposase-Accessible Chromatin using sequencing (ATAC-seq) and mutational state by whole exome sequencing (WES). Given the close link between metabolism and epigenetic regulation (*9,21*), we were interested in determining whether oHSCs with different levels of autophagy engagement could represent a set of HSC clones with distinct epigenetic features or mutagenic burden.

**Fig. 1.**
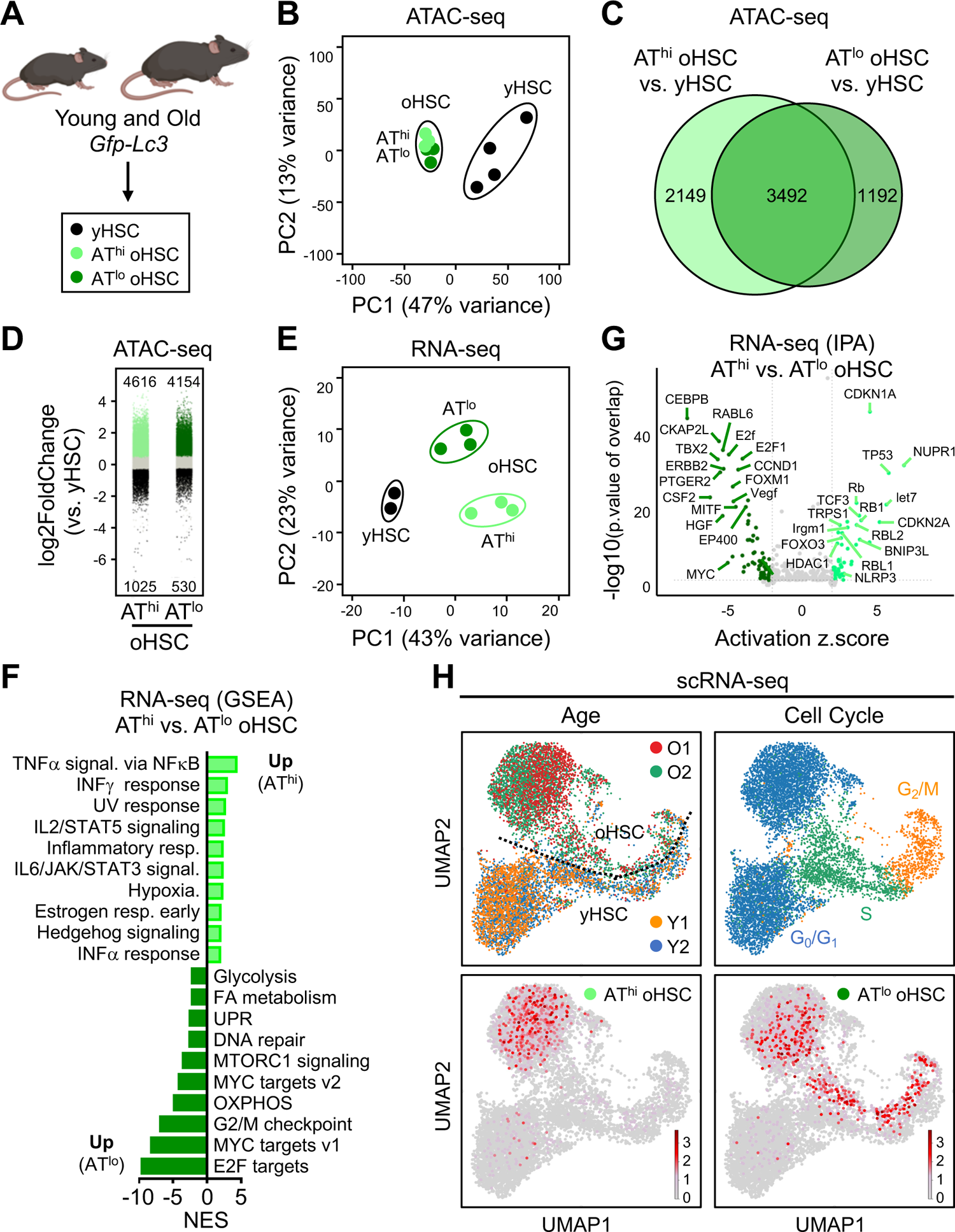
Autophagy activation in old HSCs is associated with quiescence maintenance despite a molecular inflammation response signature. (A) HSC isolation strategy from young (2-3 months) and old *Gfp-Lc3* (24-28 months) mice. (B) Principal component (PC) analysis of ATAC-seq peak counts at all genomic loci from yHSC, AT^hi^ oHSC, and AT^lo^ oHSC. (C) Venn diagram of differentially accessible (DA) peaks (LFC >=1 or <=-1, padj < 0.05) between AT^hi^ oHSC vs. yHSC and AT^lo^ oHSC vs. yHSC. (D) Directionality of DA peaks between AT^hi^ and AT^lo^ oHSC vs. yHSC. (E) PC analysis of RNA-seq gene counts from yHSC, AT^hi^ oHSC, and AT^lo^ oHSC. (F) Gene set enrichment analysis (GSEA) of AT^hi^ vs. AT^lo^ oHSC differentially expressed (DE) genes visualizing top 10 enriched and suppressed pathways by normalized enrichment score (NES) with FDR q-val <0.05. (G) Ingenuity Pathway Analysis (IPA) upstream regulators enriched from DE genes (LFC >=1 or <=-1, padj < 0.05) in AT^hi^ vs. AT^lo^ oHSC. Enriched regulators defined as Zscore >= 2 or <= −2 and p-val of overlap <0.05. (H) Uniform manifold approximation and projection (UMAP) of yHSC and oHSC 10X Genomics scRNA-seq analyses from two biological replicates (top left) with cell cycle annotation (top right). Projection of index sorted AT^hi^ (bottom left) and AT^lo^ oHSCs (bottom right) transcriptomes from SMART-Seq2 scRNA-seq onto 10X Genomics UMAP.

By ATAC-seq, we observed large differences in chromatin accessibility between young and old HSCs (**Fig. 1B**). However, we found that the chromatin accessibility landscape of AT^hi^ and AT^lo^ oHSCs was largely conserved, without statistically differentially accessible peaks between AT^hi^ and AT^lo^ oHSCs, and predominantly shared differentially accessible peaks between the two oHSC subsets and yHSC (**Fig. 1C**). Both oHSC subsets had a unidirectional increase in peak accessibility across greater than four thousand statistically significant loci compared to yHSCs, indicating epigenetic de-repression with aging (**Fig. 1D; fig. S1c**). Pathway analysis of ATAC-seq peaks differing between oHSCs and yHSCs revealed a common epigenetic poising towards engagement of inflammatory responses, cell stress, and altered metabolic programs in both AT^hi^ and AT^lo^ oHSC subsets, with some of the most differentially accessible peaks located in the promoters of genes involved in lipid metabolism (**fig. S1C,D**). Motif analysis inferred differential activity of transcriptional regulators involved in these processes such as FoxO1, Sp1/2/3, and Ikzf1 (**fig. S2a**). These results suggest that cellular responses associated with oHSC dysfunction are, to some extent, embedded in their epigenetic state. They also indicate that AT^hi^ oHSCs are similarly epigenetically poised as AT^lo^ oHSCs, and thus unlikely to represent oHSCs with a different life history. To complement these findings, we used WES to call somatic mutations in AT^hi^ and AT^lo^ oHSC subsets relative to internal tail controls (**fig. S2B,C**). Few mutations were called at a variant allele frequency of ≥ 5% in this bulk sequencing approach, in line with recent results obtained with more sensitive methods (*22*). We also observed a high degree of shared mutations between AT^hi^ and AT^lo^ oHSCs within each biological replicate, and no conserved mutations across biological replicates. The majority of called mutations were intronic or synonymous single nucleotide variants with limited expected consequences for their encoded proteins. Taken together, these results indicate that differing autophagy levels in oHSCs are not explained by clonal divergence or differential mutagenic burden, nor do they point to mutagenic burden as a driver of oHSC dysfunction in 24-month-old mice, instead suggesting that differences in autophagy status might be environmentally mediated.

### Molecular inflammatory response in autophagy-engaged quiescent old HSCs

Next, we performed detailed transcriptomic analyses of AT^hi^ and AT^lo^ oHSCs compared to yHSCs using both bulk and single cell RNA sequencing (RNA-seq). Despite similar chromatin landscape, molecular signatures from bulk RNA-seq analyses indicated a large transcriptional divergence between the two oHSC subsets (**Fig. 1E**), with pathway analyses of differentially expressed genes (DEG) demonstrating enrichment in inflammatory signaling in AT^hi^ oHSCs and in oxidative metabolism signaling in AT^lo^ oHSCs (**Fig. 1F**). Ingenuity Pathway Analysis (IPA) inferred that several inflammation-coupled pro-autophagy regulators including p53, Bnip3l, Nupr1, and Nlrp3 (*23,24*), in addition to quiescence enforcing checkpoints FoxO3a, Rb1, and Cdkn1a, were activated in AT^hi^ oHSCs (**Fig. 1G**). To gain additional resolution into the transcriptional wiring of aged HSCs, we index-sorted AT^hi^ and AT^lo^ oHSCs for plate-based SmartSeq-2 single cell RNA-seq (scRNA-seq) analyses along with unfractionated oHSC and yHSC for droplet-based 10X Genomics scRNA-seq (**Fig. 1H**). 10X Genomics data harmonized by nearest neighbor integration and uniform manifold approximation and projection (UMAP) representation distinguished yHSC from oHSC, with the largest transcriptional differences observed in the G_0_/G_1_ cell cycle phase cluster. Within oHSCs, AT^hi^ oHSCs were almost exclusively observed in the G_0_/G_1_ cluster, whereas AT^lo^ oHSCs were spread across the activation continuum, with cells still in the G_0_/G_1_ cluster found more proximal to S cluster cells (**Fig. 1H**). SmartSeq-2 scRNA-seq analyses confirmed the spread of yHSCs along the activation continuum and showed intermediate features in oHSCs with intermediate autophagy levels (AT^mid^, 33% of cells with middle GFP levels) (**fig. S2D**). Finally, most changes in the transcriptome of oHSC vs. yHSC G_0_/G_1_ cluster cells analyzed by droplet-based 10X Genomics were differences in activation mechanisms and metabolic pathways (**fig. S2E**), which mainly reflected the molecular identity of AT^lo^ oHSCs and illustrated the difficulty of identifying a minority signature like inflammation in AT^hi^ oHSCs with low resolution scRNA-seq approaches. Collectively, these results uncovered an epigenetic predisposition towards metabolic activation and cellular stress response signaling in aged HSCs, which could reflect either memory of prior stimuli (*25,26*) or transcriptional poising (*27*). We hypothesize that diversity in autophagy flux could reflect the differential engagement of an inflammation-associated quiescence-enforcing gene program, likely activated in AT^hi^ oHSCs in response to the inflamed old BM niche (*5*).

### Acute inflammation transiently activates autophagy in young HSCs

To first establish whether autophagy could be directly activated in HSCs by pro-inflammatory signals, we injected young *Gfp-Lc3* reporter mice with IFNγ or TNFα recombinant cytokines, the two most enriched inflammatory signaling pathways observed in AT^hi^ oHSCs (**Fig. 1F**), using previously established *in vivo* models of acute inflammation (*28,29*). Following a single injection of 10 µg IFNγ, we observed a strong right-shift in yHSC GFP-LC3 fluorescence at 3 hours indicative of autophagy repression and biosynthetic activation, followed at 48 hours by a significant left-shift indicating autophagy engagement, which normalized to basal levels by 96 hours and immediately preceded the return to quiescence of IFNγ-challenged HSCs (**Fig. 2A,B**). Similar autophagy induction was also observed at 48 hours post-TNFα treatment (3 injections of 2 µg TNFα every 12 hours) coinciding with when TNFα-challenged HSCs returned to quiescence (**fig. S3A**) (*29*). We then assessed whether autophagy was genetically required to prevent HSC apoptosis in response to acute inflammation as we had previously shown for metabolic stress (*3*). At 96 hours following IFNγ injection, when we expected HSCs to have engaged autophagy and returned to quiescence, we found significantly depleted HSC numbers in an established autophagy-deficient *Atg12^fl/fl^*:*Mx1*-Cre (*Atg12^cKO^*) mouse model (*3*) as compared to control (Ctrl) mice (**Fig. 2C**), suggestive of cell death absent autophagy. To confirm the cytoprotective effect of autophagy engagement, we isolated *Atg12^cKO^* and Ctrl HSCs and assessed cell expansion and caspase activation in liquid culture ± 1 mg/ml IFNγ. While IFNγ had a restraining effect on Ctrl HSC proliferation, it completely abrogated cell expansion in *Atg12^cKO^* HSCs due to apoptosis (**Fig. 2D**). We also tested the effect of TNFα injection *in vivo* but observed significant death of TNFα-treated *Atg12^cKO^* mice that precluded detailed analysis of HSC cellularity (**fig. S3B**). However, similarly reduced cell expansion and increased cell death were observed *ex vivo* upon treatment of *Atg12^cKO^* HSCs with 1 µg/ml TNFα (**fig. S3C**). Collectively, these results demonstrate a requirement for autophagy engagement in quiescence enforcement and survival in response to acute inflammatory challenge.

**Fig. 2.**
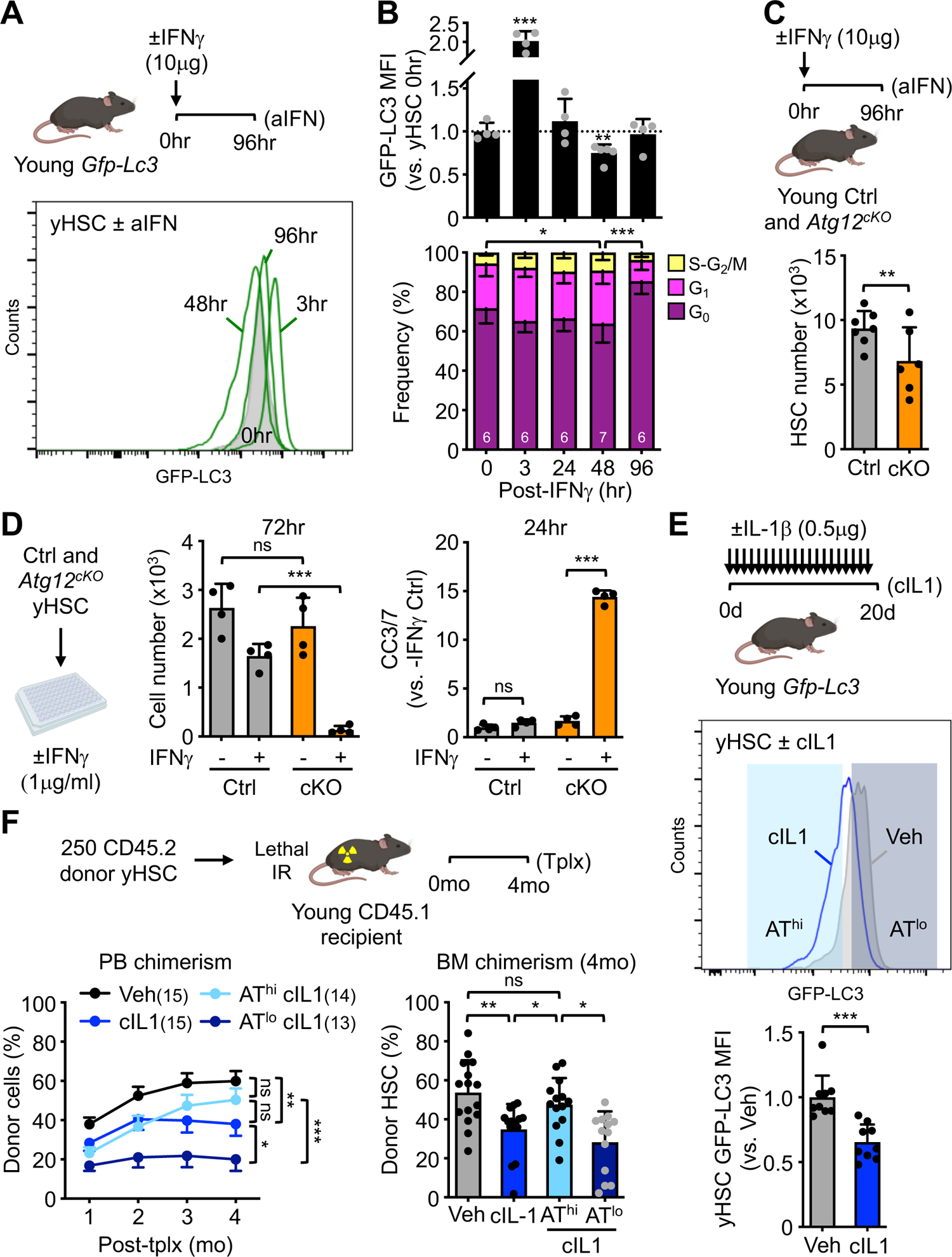
Inflammatory stimuli activate a protective autophagy program in HSC. (**A**) Experimental scheme for acute IFNγ treatment (aIFN) in *Gfp-Lc3* mice (top) with representative FACS plot of GFP-LC3 levels in HSCs (bottom). (**B**) Normalized GFP-LC3 mean fluorescence intensity (MFI, top) and Dapi/Ki67 cell cycle status (bottom) at the indicated timepoints following IFNγ injection. (**C**) HSC cellularity at 96hr following IFNγ injection in *Atg12^cKO^* (cKO) and control (Ctrl) mice. (**D**) Changes in cell number and cleaved caspase 3/7 (CC3/7) levels in *Atg12^cKO^* and control HSCs treated with or without (±) 1μg/ml IFNγ. (**E**) Experimental scheme for chronic IL-1ϕ3 treatment (cIL1) in *Gfp-Lc3* mice (top) with representative FACS plot of GFP-LC3 levels (middle) and quantification of GFP-LC3 MFI (bottom) in HSCs. (**F**) Regenerative capacity of the indicated HSC populations following transplantation (Tplx) into lethally irradiated (IR) recipients. Results show overall engraftment in the peripheral blood (PB) over time (left) and HSC chimerism in donor bone marrow (BM) at 4 months (mo) post-Tplx. Data are means ± S.D. except for overall engraftment data (F) (± S.E.M.); *p ≤ 0.05, **p ≤ 0.01, ***p ≤ 0.001; *ns*, not significant.

### Chronic inflammation promotes basal autophagy engagement to protect HSC function

We then extended these analyses to a model of chronic inflammation, where young *Gfp-Lc3* mice received daily injections of 0.5 µg recombinant IL-1β (cIL1) or vehicle (Veh) for 20 days (**Fig. 2E**), which mimics important aspects of hematopoietic and BM niche aging, including compromised HSC lineage and regenerative potential (*30*), and stromal inflammation (*5*). We found pronounced basal autophagy activation in cIL1-exposed yHSCs, from which we isolated autophagy-activated (AT^hi^, 33% low GFP) and autophagy-inactivated (AT^lo^, 33% high GFP) subsets (**Fig. 2E**). While AT^hi^ cIL1-exposed yHSCs maintained high regenerative output and HSC BM chimerism at 4 months post-transplantation, like Veh-treated yHSCs, AT^lo^ cIL1-exposed yHSCs displayed even more impaired reconstitution potential than bulk cIL-1-exposed yHSCs and significantly decreased HSC BM chimerism compared to Veh-treated yHSCs (**Fig. 2F**). Remarkably, the engraftment pattern of AT^hi^ and AT^lo^ cIL1-exposed yHSC mirrored those of AT^hi^ and AT^lo^ oHSCs (*4*), with engagement of autophagy associated in both cases with maintenance of HSC regenerative potential. However, in contrast to aging, none of the cIL1-exposed yHSC subsets displayed altered lineage distribution at 4 months post-transplantation compared to Veh-treated yHSCs (**fig. S3D**). In addition, while cIL1 exposure induced a pleiotropic inflammatory response in BM fluids (**fig. S3E,F**), it appeared significantly distinct from physiological aging (*5*) despite some shared increases in pro-inflammatory cytokines like IL-1β itself. Taken together, these results demonstrate the protective effect of autophagy on HSC regenerative capacity in response to age-related chronic inflammation. They also suggest that oHSCs could toggle between an autophagy-activated, quiescence-protected state and a metabolically activated, autophagy-repressed state based on changes in environmental stimuli, which would explain the reversibility of AT^hi^ to AT^lo^ state observed upon serial transplantation *in vivo*, and the ability of both subsets to engage autophagy upon cytokine withdrawal *in vitro* (*4*).

### Chronic inflammation impairs AKT signaling and glycolytic activity in old HSCs

To understand the metabolic consequences of chronic inflammation for HSC function, we first investigated the status of the PI3K/AKT/FoxO signaling cascade in both cIL1-exposed yHSCs and oHSCs (**Fig. 3A**). Phospho-Flow analyses showed significantly reduced AKT phosphorylation at threonine 308 (T308) and pFoxO1/3 at threonine 24 (T24) in cIL1-exposed yHSCs, with oHSCs mainly displaying reduced phosphorylation of FoxO1/3 proteins at both T24 and serine 256 (S256) sites downstream of AKT signaling consistent with FoxO activation and autophagy engagement (*31*). To further demonstrate impaired AKT signaling in aging HSCs, we directly tested the ability of oHSCs to phosphorylate AKT at threonine 308 (T308) and serine 473 (S473) in response to acute *ex vivo* treatment with 100 mg/ml insulin-like growth factor 1 (IGF-1) (**fig. S4A**). While yHSCs displayed the expected increase in pAKT^T308^ and pAKT^S473^ levels following IGF-1 stimulation, oHSCs showed no changes in AKT phosphorylation upon IGF-1 stimulation, which confirmed the defective engagement of the nutrient sensing cascade in old HSCs beyond the already reported middle-aged period (*32*). We also showed that oHSCs had decreased glucose uptake upon *in vivo* injection of a fluorescent glucose analogue (2-NBDG), indicating impaired steady state glucose metabolism, and confirmed reduced glycolysis rates by Seahorse assays (*4*), indicating impaired capacity for glycolytic activation under regenerative stimuli (**Fig. 3B,C; fig. S4B**). We also found decreased surface expression of the glucose transporter Glut1 in oHSCs compared to yHSCs, despite similar intracellular stores (**Fig. 3D; fig. S4B**). Similar features were observed in cIL1-exposed yHSCs, with scRNA-seq analyses showing enrichment of a broad inflammation response program and decreased capacity to activate demand-induced metabolism in HSCs, re-analysis of a published bulk RNA-seq dataset (*33*) indicating a predominant signature of downregulated insulin/IGF-1 pathway signaling, and Seahorse assays demonstrating markedly impaired glycolytic capacity associated with decreased glucose uptake *ex vivo* (**Fig. 3E; fig. S4C-F**). Together, these results demonstrate impaired ability to activate glycolysis and reduced steady state glucose uptake as a consequence of chronic inflammation in aging HSCs.

**Fig. 3.**
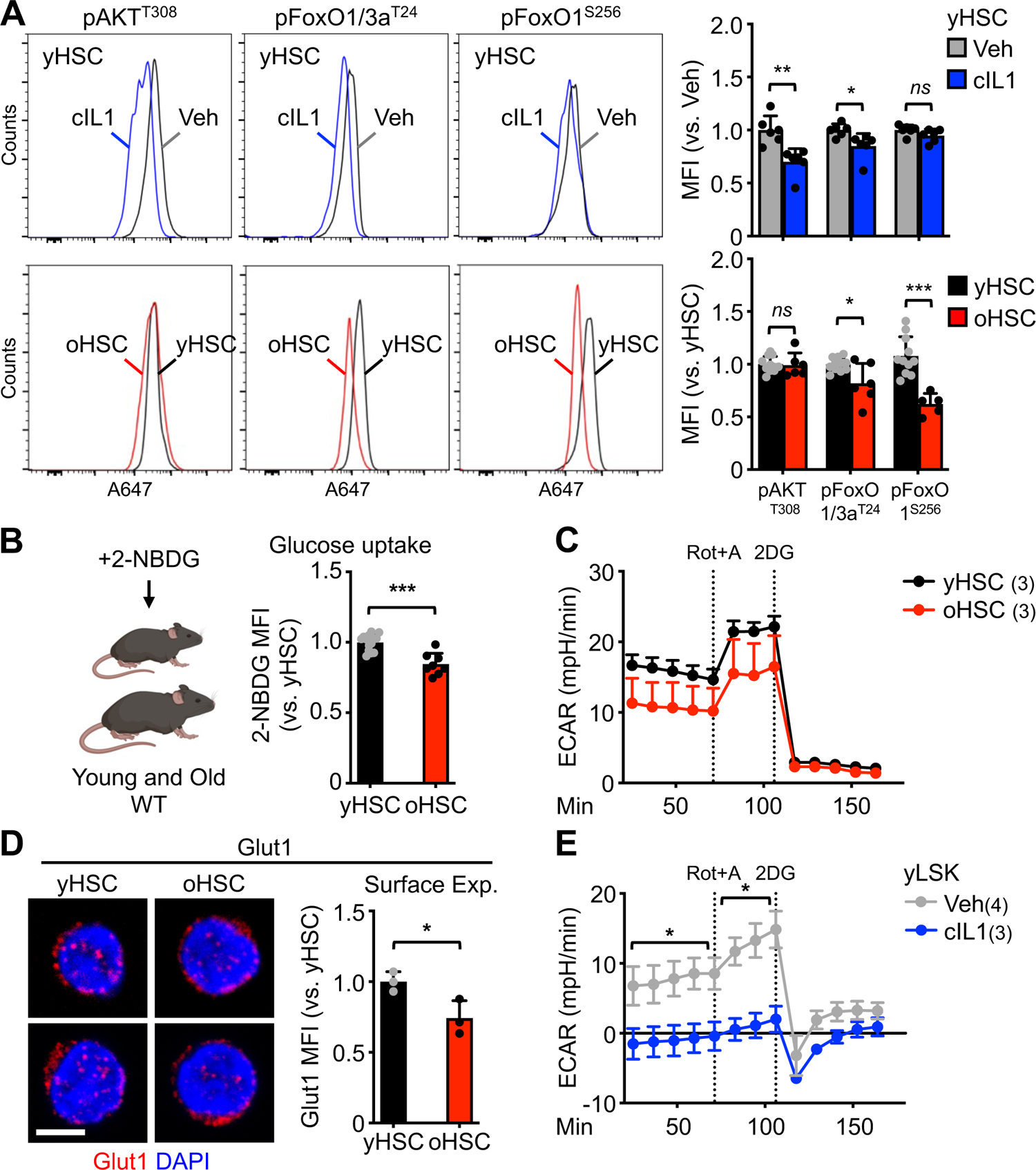
Chronic inflammatory suppresses PI3K/AKT/FoxO signaling and impairs glycolytic activity in HSCs. (**A**) Representative FACS plots and quantification of pAkt (T308), pFoxO1/3a (T24), and pFoxO1 (S256) MFI in vehicle (Veh) or chronic IL-1β (cIL1) treated young HSCs (top), and young and old HSCs (bottom). (**B**) *In vivo* 2-NBDG uptake in young and old HSCs. (**C**) Extracellular acidification rates (ECAR) measured by Seahorse glycolytic rate assay in yHSCs and oHSCs; Rot+A, rotenone + antimycin; 2DG: 2-deoxy-glucose. (**D**) Representative immunofluorescent image of Glut1 expression (left; scale bar, 10 µm) and quantification of Glut1 surface expression by flow cytometry (right) in young and old HSCs. (**E**) ECAR measured by Seahorse glycolytic rate assay in Veh and cIL1 HSC-enriched Lin^-^/c-Kit^+^/Sca-1^+^ (LSK) BM cells. Data are means ± S.D. except for Seahorse results (C,E) (± S.E.M.); *p σ; 0.05, **p σ; 0.01, ***p σ; 0.001; *ns*, not significant.

### Socs3 upregulation mediates the metabolic consequences of inflammation

Interestingly, one of the most differentially expressed genes between young and old HSCs, which was also most prominently upregulated in AT^hi^ HSCs as well as yHSCs exposed to both acute and chronic inflammatory challenges, was the suppressor of cytokine signaling 3 (*Socs3*) gene (**Fig. 4A**; **fig. S5A**).

**Fig. 4.**
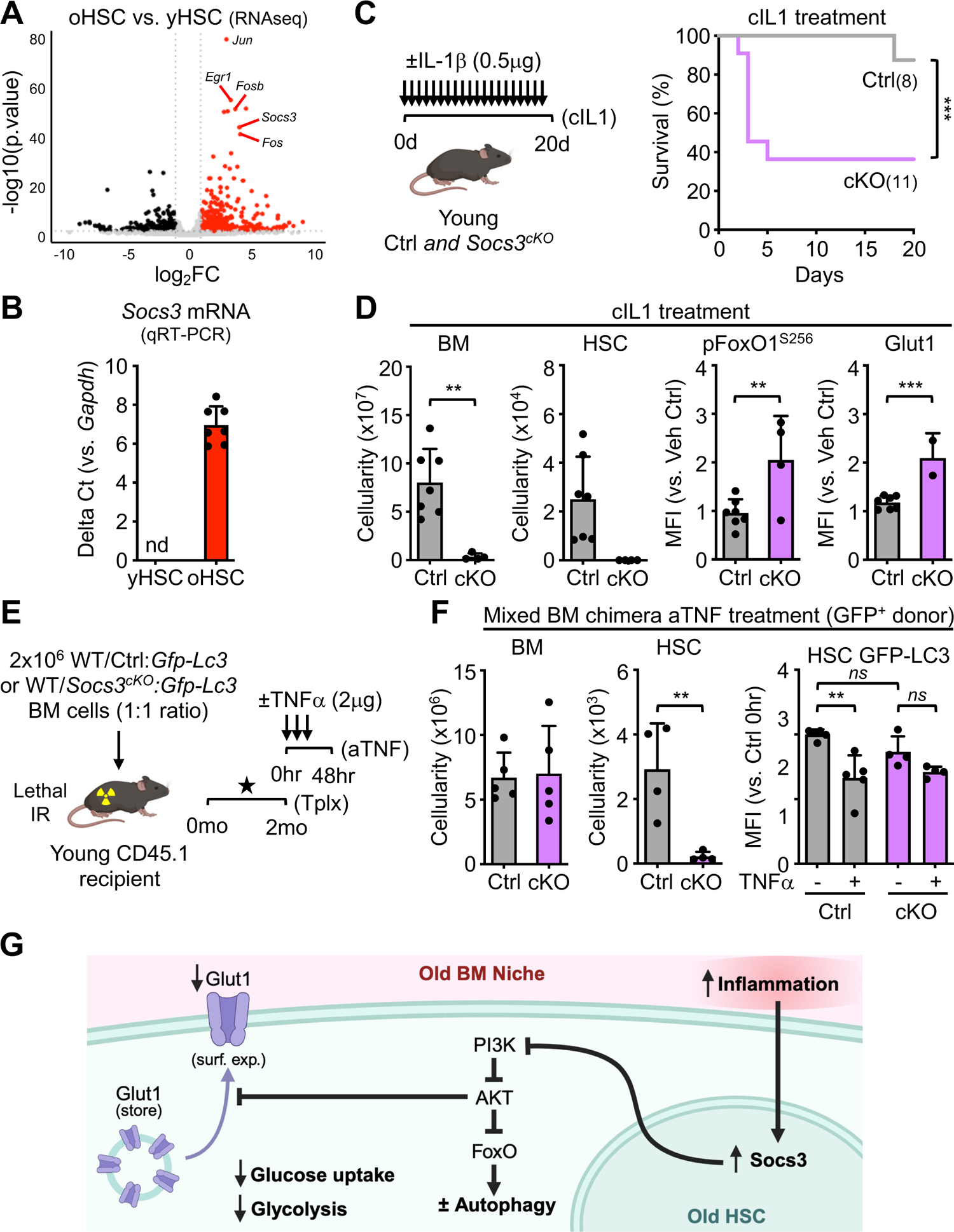
Socs3 mediates the effect of chronic inflammation on HSC function. (**A**) Volcano plot of DE genes in oHSC vs. yHSC bulk RNA-seq highlighting *Socs3* and AP-1 genes upregulation in oHSCs. (**B**) qRT-PCR confirmation of *Socs3* expression in oHSCs; nd, not detected. (**C**) Survival of Ctrl and *Socs3^cKO^*mice following 20 days of chronic IL-1β (cIL1) treatment. (**D**) Characterization of cIL1-exposed Ctrl and *Socs3^cKO^* mice for BM and HSC cellularity, pFoxO1^S256^ levels and Glut1 surface expression. (**E**) Schematic of the mixed Ctrl and *Socs3^cKO^* BM chimera used for acute TNFα (aTNF) treatment. Lethally irradiated (IR) recipients were reconstituted with the indicated 1:1 mix of wild type (WT) competitor and GFP^+^ donor BM cells, treated with poly(IC) to induce Socs3 deletion at 1 month (star), and exposed to TNFα at 2 months post-transplantation (Tplx). (**F**) Characterization of aTNF-exposed donor Ctrl and *Socs3^cKO^* cells for BM and HSC cellularity, and autophagy engagement by normalized GFP-LC3 mean fluorescence intensity (MFI). (**G**) Proposed model for the impact of chronic old BM niche inflammation on the AKT-FoxO phosphorylation cascade with consequences for glucose uptake, glycolytic capacity, and autophagy induction in old HSCs. Data are means ± S.D.; *p ≤ 0.05, **p ≤ 0.01, ***p ≤ 0.001; *ns*, not significant.

Socs3 is a well-known inflammatory mediator and inhibitor of JAK/STAT and insulin/IGF-1 signaling pathways in many tissues (*34,35*). We confirmed the robustly increased expression of *Socs3* in oHSCs by qRT-PCR (**Fig. 4B**). To directly test the role of Socs3 as a mediator of metabolic adaptations under inflammatory conditions, we generated *Socs3^fl/fl^:Mx1-Cre (Socs3^cKO^*) (*36*) to conditionally inactivate *Socs3* in HSCs upon poly(IC) treatment, and also crossed *Socs3^cKO^* mice with *Gfp-Lc3* reporter mice to monitor autophagy levels. While no adverse effects were observed upon poly(IC) injection in either *Socs3^cKO^* or *Socs3^fl/fl^*littermate control (Ctrl) mice, injection one month later with any pro-inflammatory cytokines caused a significant lethality in *Socs3*-deficient mice. For instance, no *Socs3^cKO^* mice survived acute TNFα challenge beyond 48 hours and over 60% of *Socs3^cKO^* mice quickly succumbed to chronic IL-1β treatment (**Fig. 4C; fig. S5B**). Moreover, cIL1-exposure lead to a major BM attrition in *Socs3^cKO^*mice a with significant loss of HSCs and remaining evidence of emergency myelopoiesis overactivation (*37*) with MPP3 and GMP amplification, and MPP4 depletion compared to the normal response observed in Ctrl mice (**Fig. 4D; fig. S5C**). However, in the remaining cIL1-exposed *Socs3^cKO^* HSCs, we observed increased pFoxO1^S256^ levels and Glut1 surface expression (**Fig. 4D**), demonstrating hyperactivation of the PI3K/AKT/FoxO pathway in response to inflammation in the absence of Socs3. To assess the ability of *Socs3^cKO^*HSCs to engage autophagy in mice capable of surviving an inflammatory challenge, we generated 1:1 mixed BM chimera with WT competitor and either Ctrl:*Gfp-Lc3* or *Socs3^cKO^*:*Gfp-Lc3* donor cells, which were treated with acute TNFα challenge 2 months post-transplantation and 1 month after poly(IC) injections to inactivate Socs3 (**Fig. 4E**). With mixed BM chimera mice now capable of withstanding TNFα exposure, we found a specific attrition of *Socs3^cKO^* HSCs accompanied with a lack of engagement of autophagy compared to the normal autophagy activation observed in surviving Ctrl HSCs after 48 hours of TNFα exposure (**Fig. 4F; fig. S5D**). These results demonstrate the inability of *Soc*s3-deficient HSCs to dampen PI3K/AKT signaling in response to inflammation with consequences to survival, autophagy induction and Glut1 surface expression. Collectively, they support a model in which chronic inflammation in the old BM niche impairs the AKT-FoxO phosphorylation cascade through Socs3 upregulation in old HSCs, which contributes to reduced glucose uptake, decreased glycolytic capacity, and impaired functionality in the entire HSC compartment (**Fig. 4G**). In this context, activation of autophagy acts as a compensatory response in oHSCs with the highest Socs3 expression (**fig. S5A**), to maintain functional quiescence and preserve some regenerative potential.

### Autophagy-engagement allows metabolic compensation in old HSCs

We next tried to assess how autophagy engagement could compensate for glycolytic impairment in oHSCs. Analysis of bulk RNA-seq datasets uncovered upregulation of many transcriptional regulators and metabolic enzymes associated with nutrient catabolism, including fatty acid (FA) oxidation and glutamine uptake in AT^hi^ oHSCs compared to AT^lo^ oHSCs and yHSCs (**fig. S6A**). To investigate metabolic adaptation, and in the absence of an established method sensitive enough to trace glucose usage in HSCs, we performed low-input shotgun metabolomics (*38,39*) on 10,000 isolated yHSCs, AT^hi^ oHSCs, and AT^lo^ oHSCs (**fig. S6B**). We observed minimal changes in metabolite abundance between the 3 HSC populations using a log_2_FC > 0.6 and p < 0.10 exploratory threshold. Compared to both oHSC subsets, yHSC showed higher levels of the glucose dependent non-oxidative pentose pathway intermediates D-sedoheptulose-7-phosphate and L-aspartic acid, which contribute to nucleotide and protein synthesis, respectively (*40*). In contrast, compared to yHSCs, both oHSC subsets had increased abundance of carnitine-containing molecules and β-oxidation intermediaries (*41*). Finally, in AT^hi^ oHSCs, we found a specific increase in the abundance of lyso-PE (16:0), an exogenous substrate of lysophospholipid metabolism, SM(d18:1), a substrate of sphingolipid metabolism, and glycero-3-phosphocholine, which promotes lipid metabolism along the Kennedy pathway and supports autophagosome biogenesis (*42*).

Targeted metabolomics also showed minimal changes between the 3 HSC populations other than higher abundance of acetyl-CoA in the more proliferative AT^lo^ oHSC subset (**fig. S6C**). To complement these analyses, we also used a CyTOF-based metabolic-enzyme and signaling pathway proteomics (*43*) on yHSCs, AT^hi^ oHSCs, and AT^lo^ oHSCs (**fig. S6D; table S1**). Consistently, both oHSC subsets showed a distribution in cell density away from proteins associated with glycolysis and mitochondrial quality control found in yHSCs, and towards those associated with lipid metabolism and oxidative phosphorylation in AT^hi^ and AT^lo^ oHSCs. Finally, since one of the most differentially expressed genes between AT^hi^ and AT^lo^ oHSC was *Ppargc1a* (**fig. S7A-C**), which encodes the master regulator of mitochondrial biogenesis and energy metabolism PGC-1α (*44*), we also generated *Ppargc1a ^fl/fl^:Mx1-Cre (Ppargc1a^cKO^*) mice (*45*) to conditionally inactivate PGC-1α in HSCs (**fig. S7D**). Despite its known regulation by the PI3K/AKT/FoxO pathway and role in facilitating lysosomal biogenesis and FA catabolism upon autophagy engagement (*44,46*), we could not demonstrate a role for PGC-1α in the mechanisms regulating metabolic compensation in old HSCs (**fig. S7E-H; Supplementary Text**), which suggest a high degree of functional redundancy. Together, these results suggest that autophagy engagement enables metabolic compensation for inflammation-driven impaired glucose metabolism and decreased glycolytic capacity in aging, perhaps mediated by a shift towards adaptive lipid metabolism (*8,9,47*), to support the quiescence and regenerative potential of AT^hi^ oHSCs.

### Fasting/refeeding resets the metabolism and functionally restores old HSCs

Lastly, we explored strategies by which autophagy could be manipulated to restore glycolytic metabolism in oHSCs. We turned to a 24-hour fasting/refeeding (F/R) paradigm, which transiently activates autophagy across many tissues in mice, and has previously shown potent effects on systemic metabolism, inflammation, immune function, and HSC resilience to chemotherapy induced toxicity (*48,49*). We found that both yHSCs and oHSCs potently engaged autophagy in the fasting phase and repressed autophagy in the refeeding phase without any overall changes in HSC numbers (**Fig. 5A,B**). Interestingly, the autophagy level in F/R oHSCs was comparable to untreated Ad libitum fed (AL) yHSCs (**Fig. 5B**), suggesting a reduced requirement for basal autophagy in oHSCs following this intervention. Remarkably, F/R, but not fasting alone, promoted a large increase in regenerative output of oHSCs following transplantation, restoring it to the engraftment level of yHSCs, which themselves maintained equivalent functionality across interventions (**Fig. 5C**). In addition, F/R oHSCs displayed reduced myeloid-biased and improved BM HSC donor chimerism (**fig. S8A**). To complement the fasting results, we also used HSCs from mice carrying a knock in (KI) mutation in the autophagy regulatory protein Beclin 1 (*Becn1^F121A/F121A^*), which leads to increased basal autophagy flux and extends organismal longevity (*50*). However, we found the same age-related increase in HSC numbers and defects in regenerative capacity in both KI mutant and age-matched Ctrl mice (**fig. S8B-D**). Similarly, oHSCs from 18-month-old WT mice fed for 3 months with chow containing the mTORC1 inhibitor and autophagy activator rapamycin did not show improved regenerative capacity compared to oHSCs from age-matched 21-month-old Ctrl mice **(fig. S8E-G**). Furthermore, oHSCs from rapamycin-fed animals also displayed similar TMRE levels and *ex vivo* glucose uptake compared to oHSCs from Ctrl mice (**fig. S8H**). Together, these results show that while autophagy plays an important role in sustaining the residual regenerative output of oHSC, promoting autophagy engagement across the entire oHSC compartment is insufficient by itself to restore the functionality of aged HSCs, perhaps because oHSCs still exist in a suboptimal metabolic state or are not optimally capable of engaging demand-activated metabolism.

**Fig. 5.**
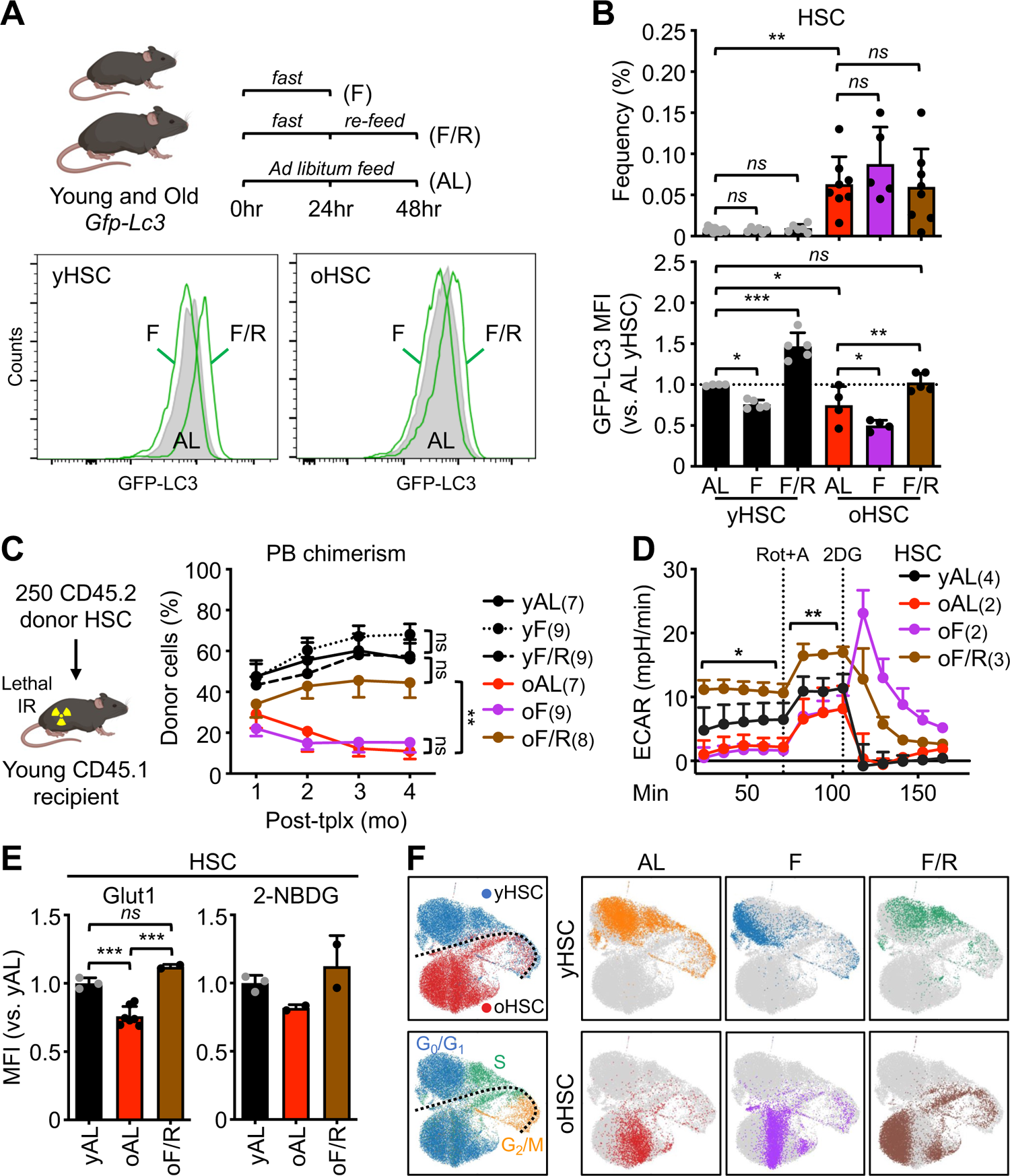
Fasting refeeding restores glycolytic and regenerative capacity of old HSCs. (**A**) Schematic of fasting and refeeding intervals in young and old *Gfp-Lc3* mice (top) with representative FACS plots of GFP-LC3 levels in HSCs in different feeding conditions (bottom). (**B**) Quantification of HSC frequency (top) and GFP-LC3 MFI (bottom) in ad libitum (AL), fasted (F) and fast/refed (F/R) states. (**C**) Regenerative capacity of the indicated young and old HSC populations following transplantation (Tplx) into lethally irradiated (IR) recipients. Results show overall engraftment in PB over time. (**D**) Extracellular acidification rates (ECAR) as measured by Seahorse glycolytic rate assay in the indicated young and old HSC populations. Rot+A, rotenone + antimycin; 2DG: 2-deoxy-glucose. (**E**) Glut1 surface expression (left) and 2-NBDG glucose uptake in the indicated young and old HSC populations. (**F**) UMAP of 10X Genomics scRNA-seq analyses of yHSC and oHSC from the indicated feeding conditions with cell cycle annotation. Data are means ± S.D. except for overall engraftment data (C) and Seahorse results (D) (± S.E.M.); *p ≤ 0.05, **p ≤ 0.01, ***p ≤ 0.001; *ns*, not significant.

To probe the metabolic basis of enhanced regenerative potential post-refeeding, we performed Seahorse analyses on HSCs isolated from AL-fed, fasted and F/R young and old mice, and found that F/R, but not fasting alone, elevated basal and maximal glycolytic capacity in oHSCs even above the levels found in yHSCs (**Fig. 5D**). In addition, we observed restoration of Glut1 surface expression and 2-NBDG glucose uptake in F/R oHSCs (**Fig. 5E**), with CyTOF analyses confirming increased Glut1 expression in F/R oHSCs upon culture activation (**fig. S9A**). These results implied that F/R intervention could restore both metabolic features and functionality of oHSCs by rescuing age-associated defects in the cellular signaling processes that regulate glycolysis. To explore the molecular basis of this phenotype, we performed scRNA-seq on AL-fed, fasted and F/R young and old HSCs (**Fig. 5F**). As previously observed, young and old HSCs were separated in the G_0_/G_1_ phase and converged upon cell cycle entry. In yHSCs, AL and F/R groups showed overlapping transcriptional signatures with a distinct pattern for the fasted group. In contrast, in oHSCs, AL and fasted groups showed a strong overlap with a distinct transcriptional signature for the F/R group, reflective of differences in metabolic set points with age. We then focused exclusively on G_0_/G_1_ cells and performed a linear regression across samples including age and feeding pattern as covariate terms, and used nearest neighbor integration to generate a UMAP, define clusters and apply a previously established HSC score (*51*) (**fig. S9B,C**). In these conditions, we observed that F/R oHSCs more closely resemble yHSCs, with a reduction in cluster 0 (c0) cells with the lowest HSC score, and increased representation of cluster 2 (c2) cells with the highest HSC score. Of interest, c2 cells were enriched primarily for pathways involved in inflammation, PI3K-AKT pathway activation and IL-7 signaling, whereas c0 cells were enriched for pathways primarily involved in respiratory metabolism and cell cycle activation (**fig. S9D**; **table S2**). Collectively, these data lend support to the concept that fasting induces transient metabolic stress that resets with refeeding and restores the function of signaling pathways controlling glycolysis, glucose metabolism, and regenerative potential of oHSCs.

## Discussion

Here, we systematically profiled aged HSCs to identify the signals that promote autophagy during aging, and the mechanisms by which autophagy induction could sustain the regenerative potential of old HSCs. We observed a non-clonal increase in chromatin accessibility at metabolic activation and cell stress loci in both AT^hi^/AT^lo^ oHSCs and uncovered a transcriptional inflammatory response program specifically associated with autophagy engagement in AT^hi^ oHSCs. We demonstrated that autophagy is activated secondary to both acute and chronic inflammatory signaling and is genetically required for HSC survival and quiescence maintenance in the face of inflammatory stimuli, thus establishing autophagy as an essential response of HSCs to inflammation. We further found that chronic inflammation impairs AKT-FoxO pathway signaling in a Socs3-dependent manner and suppresses both basal glucose metabolism and demand-induced glycolytic capacity in HSCs in a way that parallels aging (*4*), and that FoxO-dependent autophagy enables metabolic and functional compensation for this impaired state. Lastly, we demonstrated that a short fasting/refeeding intervention can reset the metabolism of the entire old HSC compartment, restoring both glucose uptake and glycolytic capacity upon stem cell activation, thereby enhancing the regenerative capacity of old HSCs (**fig. S10**). This adds a novel and much needed mechanism to the previous observation of the effect of prolong fasting on relieving chemotherapy-induced immunosuppression (*48*).

Our results describe two new and impactful discoveries that change our understanding of HSC dysfunction in aging. First, we show that autophagy is activated secondary to inflammation-induced metabolic deregulation in old HSCs. We identify chronic inflammation in the aged BM niche (*1,5,7,8*) as a central contributor to impaired regenerative potential of old HSCs, due to suppression of AKT pathway signaling and impairment of glucose metabolism and glycolytic capacity. We found that autophagy provides an essential metabolic compensation mechanism allowing some old HSCs to maintain functional quiescence and regenerative potential. Second, we show that the transient modulation of autophagy through a fasting/refeeding intervention can reset old HSC glycolytic and regenerative capacity. This finding is particularly important not only from a translational standpoint, but also because it reframes our thinking of what autophagy activation in aging is doing. We demonstrate that autophagy activation is, by itself, not sufficient to restore the regenerative capacity of old HSCs, but instead preserves the residual regenerative output of the aging HSC compartment and compensates for an impaired metabolic state. We found that the refeeding phase, where autophagy is turned off, restores the metabolic and regenerative capacity of old HSCs, and is linked to a reset of the receptivity to nutrient sensing activation signals through the AKT pathway in the aged BM niche. Therefore, our results provide essential characterization of the temporal regulation of autophagy-activating strategies needed for resetting the metabolism and restoring the function of old HSCs.

## Acknowledgements

We thank Drs. S. Sebti and B. Levine (UT Southwestern) for shipping us bones from *Becn1^KI^* mice, O. Olson (CUIMC) for help with Seahorse assays, D. Ho (Stanford) for assistance with CyTOF sample processing, M. Martin-Sandoval and T. Mathews (UT Southwestern) for help generating metabolomic data, M. Kissner for management of the CSCI Flow Cytometry Core facilities, and all members of the Passegué laboratory for critical insights and suggestions.

## Author contributions

Conceptualization: PVD, EP

Methodology: PVD, MAP, FJC-N, XW, BG, FH, MA, PF, SCB, AV, ZZ, SM, JWS, TTH Investigation: PVD, MAP, CAM, FH, PF, AV, JWS, TTH

Visualization: PVD, MAP, FJC-N, XW, MA, ZZ, JWS Funding acquisition: BG, EP

Project administration: EP Supervision: SCB, SM, BG, EP Writing – original draft: PVD, EP

Writing – review & editing: PVD, MAP, EP

## Data and materials availability

All data are available in the main text or the supplementary materials. Datasets used in these analyses have been deposited in the Gene Expression Omnibus under accession number GSE229137. Correspondence and requests for materials should be addressed to E.P.

Supplementary Materials

Materials and Methods

Material references (51 to 59)

Supplementary Text (fig. S7)

Figs. S1 to S10

Tables S1 and S2

