## Supplementary material for "Autophagy counters inflammation-driven glycolytic impairment in aging hematopoietic stem cells"

**The PDF file includes:**

Materials and Methods
Material references (52 to 60)
Supplementary Text (fig. S7)
Figs. S1 to S10
Tables S1 and S2

### Materials and Methods

Data reporting

No statistical methods were used to predetermine sample size. The experiments were not randomized, and investigators were not blinded to allocation during experiments and outcome assessment.

Mice

Young and old wild-type (WT) C57Bl6-CD45.2 and C57Bl/6-CD45.1 mice of both sexes were bred and aged in house at CUIMC. Some old WT C57Bl6-CD45.2 mice were also obtained from the National Institute on Aging (NIA) aged rodent colonies. *Gfp-Lc3* and *Atg12*$^{cKO}$ mice have been described previously (*4*). *Socs3*-deficient and *Ppargc1a*-deficient conditional knock out (cKO) mice were generated by crossing *Socs3*$^{fl/fl}$ (JAX #010944) (*36*) or *Ppargc1a*$^{fl/fl}$ (JAX #009666) (*44*) mice with *Mx1*-Cre mice. Colonies of *Gfp-Lc3*, *Atg12*$^{cKO}$, *Socs3*$^{cKO}$, *Ppargc1a*$^{cKO}$ mice of both sexes were bred and aged in house at CUIMC. At the time of analyses, young WT or *Gfp-Lc3* mice were ~ 2 months of age (range 6 to 12 weeks) and old mice ~ 24 months of age (range of 22 to 29 months). For Cre-mediated deletion, 4-8-week-old *Atg12*$^{cKO}$, *Socs3*$^{cKO}$, and respective littermate control (Ctrl) mice without the *Mx1-Cre* transgene were injected intraperitoneally three times 2 days apart with 125 µg poly(IC) (GE Healthcare) in 100 µl PBS. Cre-mediated deletion in *Ppargc1a*$^{cKO}$ instead required 6 injections of 250 µg poly(IC) 2 days apart, treatment that was also applied to littermate Ctrl mice. Poly(I/C) treated mice were only used 1 to 3 months after the last injection (for experiments with young mice including mixed BM chimera) or let aged until 24 months of age (for experiments with old mice) to ensure no residual inflammation from the poly(IC) treatment. Bones from young (~ 2-3 months of age) and old (~18 months of age) *Becn1*$^{F121A/F121A}$ knock-in (KI) mice (*50*) were shipped to us from the laboratory of Dr. Beth Levine at UT Southwestern (Dallas, TX). Young C57Bl/6-CD45.1 recipient mice for transplantation assays were ~ 2 months of age at time of irradiation. No specific randomization or blinding protocol was used, and both male and female animals were used indiscriminately in all the experiments. All mice were maintained in mouse facilities at CUIMC in accordance with IACUC approved protocols.

*In vivo* assays

For acute TNFα treatment, mice were injected retro-orbitally with 2 µg mouse TNFα (Genentech) in 100 µl PBS or 100 µl PBS alone (vehicle) 3 times every 12 hours. For acute IFNγ treatment, mice were injected retro-orbitally with 10 µg mouse IFNγ (Peprotech) in 100 µl 0.1%BSA/PBS or 100 µl 0.1%BSA/PBS alone (vehicle) once. For chronic IL-1 treatment, mice were injected intraperitoneally with 0.5 µg IL-1β (Peprotech) in 100 µl PBS/0.2% BSA or 100 µl PBS/0.2% BSA alone (vehicle) once daily for 20 days. For repeated 5-fluorouracil (5- FU) treatment, mice were injected intraperitoneally with 150 mg/kg 5-FU (Sigma-Aldrich, F6627) in PBS once a month for 2 months (2 times). For rapamycin treatment, 18-month-old WT mice were placed on either regular chow or rapamycin-containing chow encapsulated at 140 ppm (Rapamycin Holdings) and fed Ad Libidum for 3 months prior to analysis at 21 months of age. For fasting and refeeding experiments, animals were single-housed and placed into a new cage free of bedding between 9 am-10 am and left with water alone for 24 hours for the fasting period. For the refeeding period, regular chow was added to the cage and animals were left for another 24 hours. All experimental mice including Ad Libidum fed mice were single housed at the start of each experiment. For *in vivo* glucose uptake assays, WT mice were injected with 375 mg of 2-NBDG (ThermoFisher) in 150 µl PBS 1 hour prior to BM harvest, and flow cytometry analyses were quickly performed within 3 hours of BM harvest.

Transplantations

C57Bl/6-CD45.1 recipient mice were irradiated using an X-ray irradiator (MultiRad225, Precision X-Ray Irradiation), and purified cells isolated from single or pooled donor C57Bl6-CD45.2 mice were delivered via retro-orbital injections. For HSC transplantations assays, recipients were lethally irradiated with 11 Gy (delivered in split doses 3 hours apart) and transplanted with 250 donors HSCs delivered together with 300,000 Sca-1-depleted helper C57Bl/6-CD45.1 BM cells. For mixed BM chimera, lethally irradiated recipients were transplanted with 2x106 BM cells representing a 1:1 ratio of WT C57Bl6-CD45.2 competitor and GFP-expressing donor cells. Transplanted mice were kept on polymyxin/neomycin containing water for 4 weeks. Peripheral blood (PB) was analyzed monthly for donor-derived chimerism via retro-orbital bleeding and collected in either 4 ml of ACK (150 mM $NH_4Cl$/10 mM $KHCO_3$) buffer containing 10 mM EDTA for flow cytometry analyses or EDTA-coated tubes (Becton Dickinson) for complete blood count (CBC) analyses using a Genesis (Oxford Science) hematology system. For each cohort, the number of transplanted mice that were followed over time and the numbers of mice that were analyzed at 4 months post-transplantation for reconstitution are indicated on the figures. Recipient mice with chimerism levels ≤ 1% in the BM were excluded from the analyses.

Cell isolation flow cytometry

BM cells were obtained by crushing with a mortar and pestle leg, arm, and pelvic bones (with sternum and spines for some experiments) in staining media composed of Hanks' buffered saline solution (HBSS) containing 2% heat inactivated FBS (Cellgro B003L52). Red blood cells (RBC) were removed by lysis with ACK buffer, and single-cell suspensions of BM cells were purified on a Ficoll gradient (Histopaque 1119, Sigma-Aldrich). PB collected in ACK buffer containing 10 mM EDTA was further lysed in ACK buffer to remove contaminating RBCs. Cellularity was determined by ViCELL-XR automated cell counter (Beckman-Coulter). For HSC isolation, BM cells were first enriched for c-Kit$^+$ cells using c-Kit microbeads (Miltenyi Biotec, 130-091-224) and an AutoMACS cell separator (Miltenyi Biotec), and then incubated with a cocktail of purified rat anti-mouse lineage antibodies including CD3 (BioLegend, 100202), CD4 (eBioscience, 16-0041-82), CD5 (BioLegend, 100602), CD8 (BioLegend, 100702), CD11b (BioLegend, 101202), B220 (BioLegend, 103202), Gr-1 (eBioscience, 14-5931-85), and Ter-119 (BioLegend, 116202). For HSC analyses, BM cells were directly incubated with purified lineage antibodies. Cells were then incubated with goat anti-rat-PE-Cy5 (Invitrogen, A10691), blocked with purified rat IgG (Sigma-Aldrich), and stained with c-Kit-APC-Cy7 (BioLegend, 105826), Sca-1-BV421 (BioLegend, 108128), Flk2-PE (eBioscience, 12-1351-82), CD150-PE-Cy7 (BioLegend, 115904), and CD48-A700 (BioLegend, 103426). For donor-derived HSC BM chimerism analyses in transplanted mice, Flk2-biotin (eBioscience, 13-1351-85) followed by SA-BV605 (BioLegend, 405229; 1:400) was used together with CD45.1-PE (eBioscience, 12-0453-83) and either CD45.2-A647 (eBioscience, A14737) or CD45.2-FITC (eBioscience, 11-0454-85) congenic markers. For donor-derived PB chimerism analyses in transplanted mice, blood cells were stained with CD11b-PE-Cy7 (eBioscience, 25-0112-82), Gr-1-e450 (eBioscience, 57-5931-82), B220-APC-Cy7 (eBioscience, 47-0452-82), CD3-APC (eBioscience, 17-0032-82), and Ter-119-PE-Cy5 (eBioscience, 15-5921-83) together with CD45.1 and CD45.2 congenic markers. Cells were finally re-suspended in staining media with 1 µg/ml propidium iodide to exclude dead cells. Cell isolation was performed on a Becton Dickinson (BD) FACS ARIA II SORP using double sorting, and cell analyses were performed on a Novocyte Quanteon (Agilent). GFP-LC3 and 2-NBDG signals were detected in the FITC channel.

Dye-based cytometry

For each mitochondrial membrane potential (TMRE) and mitochondrial mass (MTG) analyses, 10 million BM cells were first stained for HSCs as described above using purified lineage antibody cocktail followed by goat anti-rat-PE-Cy5, c-Kit-APC-Cy7, Sca-1-BV421, Flk2-biotin, CD150-PE-Cy7, CD48-A700, and finally SA-BV605. Samples were then incubated for 30 min at 37 °C with 100 nM tetramethylrhodamine-ethyl-ester (Enzo, enz-52309) or 30 nM mitotracker green (ThermoFisher, M7514) together with 50 μM verapamil (Sigma Aldrich, V4629) in staining media as described (*52*). Samples were finally washed twice and resuspended in staining media containing 1 μg/ml propidium iodide, and analyzed for dye fluorescence in the PE channel for TMRE and FITC channel for MTG.

Phospho-flow cytometry

For phospho-flow analysis, BM from 1 femur per sample was flushed with 4% PFA/PBS to immediately fix the samples following animal euthanasia and processed as described (*12*). Briefly, flushed samples were fixed for 10 min at RT, spun down at 2,000 rpm for 5 min, and resuspended in the fixation buffer of the FoxP3 Transcription Factor Detection Kit (eBioscience, 00-5523-00) for 1 hour at 4°C. Samples were then washed with the permeabilization buffer from the staining kit and resuspended in the same permeabilization buffer containing a cocktail of PE-Cy5 directly conjugated lineage antibodies including CD3 (eBioscience, 15-0031-83), CD4 (eBioscience, 15-00410-82), CD5 (BioLegend, 100610), CD8 (eBioscience, 10-0081-82), B220 (eBioscience, 15-0452-82), CD11b (eBioscience, 15-0112-82), Gr-1 (eBioscience, 15-5931-82), and Ter-119 (eBioscience, 15-5921-83) together with cKit-APC-Cy7, Sca-1-BV421, Flk2-PE, CD150-PE-Cy7, CD48-A700, and 1:400 dilution of either monoclonal rabbit anti-pT308 AKT (Cell Signaling, 13038S), polyclonal rabbit anti-pS256 FoxO1 (Cell Signaling, 9461S), or polyclonal rabbit anti-pT24 FoxO1/pT32 FoxO3a (Cell Signaling, 9464S) primary antibodies for 1 hour at RT. Following washing, samples were resuspended in permeabilization buffer containing 1:400 dilution of donkey anti-rabbit A647 (Jackson ImmunoResearch, 711-605-152) secondary antibody for 30 min at RT. Samples were finally resuspended in PBS and analyzed by flow cytometry on a Novocyte Quanteon (Agilent).

Cell cycle flow cytometry

For Dapi/Ki67 analysis, unfractionated BM cells were stained with a cocktail of PE-Cy5 directly conjugated lineage antibodies (CD3, CD4, CD5, CD8, B220, CD11b, Gr-1, and Ter-119) together with cKit-APC-Cy7, Sca-1-PE-Cy7 (BioLegend, 108114), ESAM-APC (BioLegend, 136207), CD150-PE (BioLegend, 115904) and CD48-A700 in flow cytometry staining media. Samples were then fixed in FoxP3 fixation buffer (eBioscience, 00-5523-00) for 1 hour at 4°C, washed in permeabilization buffer, and stained with Ki67-FITC (eBioscience, 11-5698-80) for 30 minutes at 4°C. Samples were washed and resuspended in 1μg/ml DAPI (ThermoFisher, D1306) in staining media and analyzed by flow cytometry on a Novocyte Quanteon (Agilent).

Glut1 flow cytometry

For surface staining of glut1 on HSCs, unfractionated BM cells were stained with the same cocktail of PE-Cy5 directly conjugated lineage antibodies (CD3, CD4, CD5, CD8, B220, CD11b, Gr-1, and Ter-119) together with cKit-APC-Cy7, Sca-1-BV421, Flk2-biotin, CD150-PE, CD48-A700, and rabbit anti-Glut1 (Abcam, ab115730) in flow cytometry staining media for 30 minutes at 4°C. Samples were then washed in staining media and stained with a combination of SA-BV605 and donkey anti-rabbit A647 for 1 hour at 4°C. Cells were finally washed with staining media, re-suspended in staining media containing 1 μg/ml propidium iodide to exclude dead cells, and analyzed by flow cytometry on a Novocyte Quanteon (Agilent).

Cell culture

All cultures were performed at 37 °C in a 5% $CO_2$ water jacket incubator (ThermoFisher). For apoptosis and cell expansion assays, 1,000 or 3,00 HSCs were seeded per well of a 96-well flat-bottom plate and cultured for 24 or 72 hours, respectively, in Iscove's modified Dulbecco's media (IMDM) medium (StemCell Technology, 06200) supplemented with 5% FBS (StemCell Technology, 06200), penicillin (50 U/ml)/streptomycin (50 μg/ml), non-essential amino acids (0.1 mM), sodium pyruvate (1 mM), L-glutamine (2 mM) and 2-mercaptoethanol (50 μM), and containing the following cytokines: SCF (25 ng/ml), Flt3L (25 ng/ml), IL-11 (25 ng/ml), IL-3 (10 ng/ml), GM-CSF (10 ng/ml), Epo (4 U/ml), and Tpo (25 ng/ml) (all from Peprotech). For apoptosis assays, after 24 hours culture, 200 cells equivalent volume (40 μl) was transferred to a 384-well white luminescence plate containing another 40 μl of media, and 40 μl of Caspase-Glo 3/7 reagent (Promega, G8091) were added to each well. Plates were then shaken at 300 rpm for 1 min, incubated for 60 min at RT and read on a luminometer (Molecular Devices). Background luminescence was determined by adding instead 40 μl of media without cells and was subtracted before calculating fold changes. For autophagy induction assays, 1,000 HSCs were seeded per well of a 96-well flat-bottom plate and cultured for up to 24 hours in StemPro-34 medium (Invitrogen) supplemented with L-glutamine (2 mM) and penicillin (50 U/ml)/streptomycin (50 μg/ml) with or without (±) the above cytokines (Cyto). GFP-LC3 level was measured by flow cytometry and detected in the FITC channel. For measurement of glucose uptake in WT HSCs, 2,000 cells were cultured in + Cyto IMDM media containing 100 μM 2-(n-(7-nitrobenz-2-oxa-1,3-diazol-4-yl)amino)-2-deoxyglucose (2-NBDG, Invitrogen, N13195). Cells were then washed once in staining media, re-suspended in staining media containing 1 μg/ml PI, and analyzed for 2-NBDG fluorescence in the FITC channel. For measurement of glucose uptake in *Gfp-Lc3* HSCs, 4,000 HSCs were cultured with 100 μM 2-deoxyglucose (2-DG) for 4 hours and then processed according to the manufacturer's instructions for detection of 2-deoxy-D-glucose-6-phosphate (2DG6P) bioluminescence (Glucose Uptake-Glo Assay, Promega, J1341).

<u>Seahorse assays</u>

For Seahorse metabolic flux experiments, oxygen consumption rates (OCR) and extracellular acidification rates (ECAR) were measured using a 96-well Seahorse Bioanalyzer XF 96 according to the manufacturer's instructions (Agilent Technologies). In brief, HSCs or $Lin^-$/$Sca\text{-}1^+$/$c\text{-}Kit^+$ (LSK) BM cells (50,000 or 75,000 cells per well) were sorted into 1.4 ml Eppendorf tubes and spun down at 4,200 rpm for 5 min. Supernatant was aspirated until only 20 μl remained and resuspended with 180 μl of Seahorse glycolytic rate assay media containing 10 mM glucose, 2 mM L-glutamine, 1M NaOH, SCF (25 ng/ml), Flt3L (25 ng/ml), IL-11 (25 ng/ml), IL-3 (10 ng/ml), GM-CSF (10 ng/ml), Epo (4 U/ml), and Tpo (25 ng/ml) in Seahorse XF base medium (Agilent, 103576-100). Resuspended cells were then transferred into 96-well plates pre-coated for 3 hours with poly-Lysine (Sigma-Aldrich, P4707) and left to equilibrated for 45 min at 37 °C in a 5% $CO_2$ water jacket incubator. Data acquisitions were performed according to the manufacturer's instructions with mix and read times extended to 7 min and 4 min, respectively, and glucose included from the start of all assays.

<u>Immunofluorescence</u>

HSCs (~ 2,000/slide) were directly deposited onto poly-lysine coated slides (Sigma-Aldrich, P0425), settled down for 15 min at RT, fixed with 4% PFA for 10 min at RT, permeabilized using 0.1% TritonX100 for 2 min at RT, and blocked overnight in 1% BSA in PBS at 4°C. Cells were then incubated with rabbit anti-Glut1 (Abcam, ab115730) primary antibody for 2 hours at RT, follow by a donkey anti-rabbit A594 (Jackson ImmunoResearch, 711-585-152) secondary antibody for 1 hour at RT, before being stained with 1 μg/ml DAPI (Sigma-Aldrich, D8417) for 10 min at RT and mounted with VectaShield (Vector Laboratories, H-1000). Cells were imaged on a SP5 Leica upright confocal microscope (63x objective), and images were processed using the Volocity software (Perkin Elmer v.6.2).

Metabolomics

Metabolomics experiments were performed in the laboratory of Dr. Sean Morrison at UT Southwestern (Dallas, TX) as previously described (*38,39*). Briefly, BM cells from young and old *Gfp-Lc3* mice were rapidly isolated at 4°C and stained with a cocktail of APC directly conjugated lineage antibodies including CD2 (Tonbo, 35-0029-T100), CD3 (Tonbo, 35-0032-U100), CD5 (BioLegend, 100606), CD8 (Tonbo, 35-0081-U500), B220 (Tonbo, 35-0452-U500), Gr-1 (Tonbo, 35-5931-U500), and Ter-119 (Tonbo, 35-5921-U500) together with cKit-APC-e780 (eBioscience, 47-1171-82), Sca-1-PerCP-Cy5.5 (BioLegend, 108124), CD150-PE, CD48-A700, and DAPI for live/death cell discrimination. The cell sorter (BD FACS ARIA II) was carefully cleaned and prepared as previously described (*39*), and 10,000 HSCs were directly isolated into Eppendorf tubes containing 40µl 100% acetonitrile for a final volume of ~ 50µl 80% acetonitrile per condition. After sorting, metabolites were quickly extracted by vortexing at high speed for 1 min followed by centrifugation at 17,000 x g for 15 min at 4°C. Liquid chromatography and mass spectrometry were performed as previously described (*38*). Liquid chromatography was performed using a Vanquish Flex UHPLC (Thermo Scientific) using a ZIC-pHILIC column (2.1 x 150, 5µm, Millipore Sigma) with a binary solvent gradient. Mobile phase A was water containing 10 mM ammonium acetate, pH 9.8 with ammonium hydroxide; mobile phase B was 100% acetonitrile. Gradient separation proceeded as follows: from 0 to 15 min mobile phase B was ramped linearly from 90% to 30%; from 15 min to 18 min, mobile phase B was held at 30%; from 18 min to 19 min, mobile phase B was ramped linearly from 30% to 90%; mobile phase B was held at 90% from 19 min to 27 min to regenerate the initial chromatographic environment. Throughout the method, solvent flow rate was kept at 250 µl/min and the column temperature was maintained at 25°C. 20µl of sample was injected onto the column. All mass spectrometry data were acquired using a QExactive HF-X mass spectrometer (Thermo Scientific) using polarity-switching MS1 only acquisition. Mass spectrometer parameters used for data acquisition were described previously. Mass spectrometry data were analyzed using Trace Finder 4.1 software (Thermo Scientific), and a library of metabolites and retention times developed by the Children's Research Institute Metabolomics Facility (*38*) was used for manual peak integration.

CyTOF Metabolic Profiling

CyTOF experiments were performed in the laboratory of Dr. Sean Bendall at Stanford University (Palo Alto, CA) as previously described (*43*), with cells processed at CUIMC using cisplatin-194 (Standard BioTools Inc.) for live/dead cell discrimination and shipped to Stanford on dry ice. HSCs from young and old *Gfp-Lc3* mice (15,000-30,000 cells per sample) and c-Kit-enriched BM cells from AL, F and F/R young and old *Gfp-Lc3* mice (3 million cells per sample) were fixed immediately upon isolation, while HSCs from young and old WT mice (15,000-30,000 cells per sample) were cultured for 21 hours in full cytokine activating conditions (SCF, Flt3L, IL-11, IL-3, GM-CSF, Epo, Tpo) before fixation. Cells were fixed in 1.6% PFA/PBS for 10 min at RT, resuspended in cell staining media (PBS with 0.5% BSA, 0.02% sodium azide) with 10% DMSO (Sigma Aldrich), and then stored at -80°C until shipment. Metal-isotope labeled antibodies conjugation, palladium-based mass tag cell barcoding, surface and intracellular antibody staining, and data acquisition on a CyTOF2 mass cytometer (Standard BioTools Inc.) were performed at Stanford as previously described (*53*). CyTOF datasets were transformed using an arcsinh scale of 5 and percentile normalized using a quantile value of 0.99. Features were z-scored (mean-centered and scaled) before dimensionality reduction by Principal Component Analysis. CyTOF Analysis was performed using R version 4.1.3, and statistical testing was performed using the CytoGLMM package (*54*).

Cytokine profiling

BM fluid was flushed out from the four hind leg bones (two femurs and two tibiae) using the same 200 µl of HBSS/2% FBS in a 1 ml syringe with 26g needle. BM cells were sedimented by centrifugation at 300g for 5 min and collected supernatants were purified by an additional centrifugation at 15,300g for 10 min. BM fluids were stored at -80°C until use. For bead array analyses, 50 µl of 2x diluted BM fluid was analyzed using Mouse 20-Plex panel (ThermoFischer) on a Luminex 200 analyzer according to manufacturer's protocol. For the mouse 200-plex cytokine array, BM fluid samples were submitted to Quantibody Testing Service (Raybiotech) and diluted 4x prior to analyses.

Quantitative RT-PCR analyses

Total RNA was isolated from ~ 5,000-10,000 HSCs isolated using the RNeasy Plus Micro Kit (Qiagen, 74034). RNA was treated with DNase I and reverse-transcribed using SuperScript III kit with random hexamers (Invitrogen). Runs were performed on a 7900HT Fast Real-Time PCR System (Applied Biosystems) using SYBR Green reagents (Applied Biosystems) and the cDNA equivalent of 200 HSC per reaction. Sequences for qRT–PCR primers were: *Ppargc1a,* forward - AAGTGGTGTAGCGACCAATCG; reverse - AATGAGGGCAATCCGTCTTCA (NM_008904.2); *Actb*, forward - GACGGCCAGGTCATCACTATTG; reverse - AGGAAGGCTGGAAAAGAGCC (NM_007393); *Socs3*, forward - GTTGAGCGTCAAGACCCAGT; reverse – GGGTGGCAAAGAAAAGGAG (NM_007707.3); and *Gapdh*, forward - GGCAAATTCAACGGCACAGT; reverse – GTCTCGCTCCTGGAAGATGG (NM_001289726.2). Values were normalized to *Actb for Ppargc1a* expression and *Gapdh* for *Socs3* expression.

Bulk RNA-seq

For the yHSC, $AT^{hi}$ oHSC, and $AT^{lo}$ oHSC experiment, RNA was isolated from ~10,000 HSCs using the RNeasy Plus Micro Kit (Qiagen, 74034). RNA integrity number (RIN) was determined by Bioanalyzer (Agilent Technologies) and RNA samples with RIN > 9.0 were further processed. RNA samples were processed over two experimental "sort days" and stored at -80°C in RNA isolation buffer until library preparation. A total of 3 paired $AT^{hi}$/$AT^{lo}$ oHSC biological replicates were sequenced alongside 2 yHSC replicates. RNA was quantified and QC using an Agilent Bioanalyzer. Illumina sequencing libraries were prepared and sequenced at the NYU Genome Technology Center (New York, NY). The NuGen Ovation Trio low input RNA-seq library preparation kit was used for library preparation, with all libraries prepared at the same time to minimize batch effects. Libraries were sequenced using an Illumina HiSEQ 4000 instrument with paired end 50 base pair sequencing to a target depth of 40 million unique reads per sample. Sequencing QC was performed using FastQC and MultiQC. Paired-end reads were mapped to the mouse reference genome (GRCm38 ver104) using STAR and quantified with FeatureCounts. Gene counts were imported into a DDS object using DESeq2, removing genes with less than 100 reads summed across all included samples. The rlog transformation implemented in DESeq2 was utilized to normalize sample read counts for exploratory data analysis, including principal component analysis (PCA) and hierarchical clustering. For yHSC, $AT^{hi}$ oHSC, and $AT^{lo}$ oHSC RNA-seq samples, we created a linear model regressing out the "sort day" batch effect, design(dds) = formula (~sortBatch + group). The Benjamini-Hochberg algorithm was used for multiple testing correction, with an FDR = 0.05 significance threshold. Gene Set Enrichment Analysis (GSEA) was performed for indicated pairwise comparisons using gene symbol log2 fold change pre-ranked files using the classic enrichment statistic, gene sets with a minimum size of 15 and maximum size of 500 genes, the mouse gene symbol remapping MSigDB.v7.0 chip, and the c2.cp.recatome.v6.2.symbols curated gene database. Ingenuity pathway analysis was used in genes passing a log 2-fold change greater than or equal to 1 or less than or equal to -1 and FDR <0.05 cutoff.

Bulk ATAC-seq

A total of 3 paired $AT^{hi}/AT^{lo}$ oHSC biological replicates and 5 yHSC replicates were generated over 2 isolation and transposition days. Young and old mice were both used each day to allow for batch correction. For each replicate, pools of 3 to 4 young *Gfp-Lc3* mice and individual old *Gfp-Lc3* mouse were used to isolate ~10,000 HSCs per sample for input. At harvest, cells were lysed and immediately transposed as described (*55*), with some minor modifications. Briefly, HSCs were sorted into 400 µl staining media, washed with 1 ml PBS and spun down at 500 g for 5 min at 4°C. Liquid was carefully aspirated and pellets were resuspended in 50 µl ATAC-seq lysis buffer (ATAC-RSB [1M Tris-HC1 pH 7.4, 5M NaCl, and 1M MgCl2 in water] with 0.1% Tween 20, 0.01% digitonin and 0.1% NP40) and left on ice for 3 min. Samples were then washed with 1 ml ATAC Wash buffer (ATAC-RSB with 0.1% Tween 20), inverted to mix, spun for 10 min at 500 g at 4°C, and resuspended in 20 µl ATAC transposition buffer (Nextera TD Buffer (Illumina) with 100 nM Nextera Transposase (Illumina), 0.01% Digitonin and 0.01% Tween 20 in PBS). DNA was transposed for 30 min at 37°C, purified using the ZYMO clean and concentrator-5 kit (Zymo), eluted in 22 µl Zymo Elution buffer, then stored at -20°C until library preparation. Libraries were pre-amplified with a common Ad1 adapter and unique Ad2 adapter sequences, and additionally amplified using qPCR cycle determination with the NEBNext® High-Fidelity 2X PCR Master Mix (NEB). All libraries were prepared on the same day to limit library preparation-associated batch effects. Double sided AMPure XP bead purification kit (Beckman Coulter) was utilized to remove primer-dimers and large fragments > 1000bp. Libraries were QC using Qubit and Agilent Bioanalyzer at the CUMC Herbert Irving Comprehensive Cancer Center Molecular Pathology core. Samples were sequenced by GENEWIZ using paired end 150bp sequencing on a HiSeq400 instrument to a target depth of 100 million unique reads per sample. Sequencing quality control was performed using FastQC and MultiQC, which revealed Nextera sequencing adapter contamination that was removed using the Trim Galore Cutadapt wrapper in paired end mode, followed by confirmation of adapter removal with FastQC. BWA was used to align samples to the GRCm38 ver 104 genome. BAM files were quality controlled by looking at the distribution of reads aligning to chromosomes and the distribution of insert sizes. Peaks were called using MACS2, removing PCR duplicates. NarrowPeak files generated by MACS2 were quality controlled for distribution of reads mapping to GRCm38 genomic regions using the R CHIPseeker package and Tx.Db.mmusculus.UCSC.mm39.refGene annotation. For differential ATAC-seq, a GRanges consensus peak object was generated using ChIPQC containing non-redundant open regions present in any sample. Nucleosome free regions (<100bp fragment length) from sample BAM files were counted using FeatureCounts against the consensus peak annotation. Sample counts were imported into a dds object using DESeq2, and exploratory data analysis was performed using rlog normalized count data and PCA. For differential peak accessibility calling between groups, we used DESeq2 using the results function and the DESeq2 estimator. For pathway enrichment analysis, we annotated promoter proximal peaks within ± 1000bp of the transcription start site (TSS) using TxDb.Mmusculus.UCSC.mm39.refGene. For AThi oHSC, ATlo oHSC, and yHSC samples, we used GSEA analysis using the GSEA software and MSigDB Hallmark pathways.

Whole Exome Sequencing

A total of 5 paired $AT^{hi}/AT^{lo}$ oHSC biological replicates were sequenced alongside tail DNA from each old *Gfp-Lc3* mouse to establish baseline germline mutations. Cell pellets containing ~ 10,000 HSCs per sample were frozen dry and sent to the Memorial Sloan Kettering Cancer Center Sequencing Core Facility (New York, NY) or to Genewiz (South Plainfield, NJ). DNA was prepared for whole exome capture and sequencing by these services. $AT^{hi}$ and $AT^{lo}$ oHSC DNA samples were sequenced to 150X coverage and tail control DNA from each biological replicate was sequenced to 30X depth with paired end sequencing.

Bowtie2 was used to align Fastq files to mouse genome, and somatic mutations relative to tail control were called with MuTect2.

Droplet-based scRNA-seq (10X Genomics)

For each experiment, ~ 30,000 HSCs were sorted into 1.5 ml tubes containing 500 µl of HBSS/2% FBS and transferred to the Columbia Genome Center Single Cell Analysis Core Facility for microfluidic cell processing, library preparation and sequencing. In brief, cells were re-counted, and viability was assessed using a Countess II FL Automated Cell Counter (Thermo), and samples were processed following manufacturer's recommendations for Chromium Single Cell 3′ Library & Gel Bead Kit v2 (10X Genomics). On average ~10,000 HSCs were loaded for each sample and 1 sample was loaded per condition. Samples were sequenced using an Illumina HiSEQ 4000 instrument. QC was performed keeping cells with more than 500 genes detected, less than 100,000 total reads, and less than 20% mitochondrial reads. Downstream analyses were done using Scanpy with cells normalized to 10,000 UMIs per cell and logarithmically transformed. Highly variable genes (HVGs) were selected using the "FindVariableFeatures" method". UMAP visualizations were obtained from 50 PCA components. Cell clusters were defined using either Louvain or Leiden algorithms.

Plate-based scRNA-seq (SMART-seq2)

Samples were processed following modifications to the Smart-Seq2 protocol based in the mcSCRB-Seq protocol (*56*). Single yHSC, $AT^{hi}$ oHSC, $AT^{mid}$ oHSC, and $AT^{lo}$ oHSC were directly sorted into individual wells of a 96-well PCR plate in 2.3 µl of lysis buffer containing 0.2% Triton X-100 (Sigma-Aldrich) and 1U of Superase-In RNase Inhibitor (Ambion). Of note, information regarding expression of surface markers was recorded for each cell when sorting. Cells were frozen immediately at -80°C until further processing. After thawing on ice, 2 µl of an annealing mixture containing1 µM oligodT (IDT), 5 mM each dNTPs and a 1:6,000,000 dilution of ERCC RNA Spike-In Mix (Invitrogen) was added followed by incubation at 72°C for 3 min. Then 5.7 µl of a Reverse Transcription mix containing 3.5 U/µl of Maxima H minus retrotranscriptase (ThermoFisher), 0.88 U/µl of Superase-In RNase Inhibitor, 1.75x Maxima RT Buffer, 3.5 µM TSO (Qiagen) and 13.15% PEG 8000 (Sigma-Aldrich) was added and the mixture was incubated at 42°C for 90 min, followed by 70°C for 15 min. cDNA was further amplified by adding 40 µl of a PCR mix containing 0.03 U/µl of Terra PCR direct polymerase (Takara Bio), 1.25x Terra PCR Direct Buffer, and 0.25 µM IS PCR primer (ID). PCR was as follows: 3 min at 98°C for initial denaturation followed by 21 cycles of 15 s at 98°C, 30 s at 65°C, 4 min at 68°C. Final elongation was performed for 10 min at 72°C. Sequences of oligodT, TSO and IS PCR primers were as previously described (*57*). Following preamplification, all samples were purified using Ampure XP beads (Beckman Coulter) at a ratio of 1:0.6 with a final elution in 25 µl of EB Buffer (Qiagen). The cDNA was then quantified using the Quant-iT PicoGreen dsDNA Assay Kit (Thermo Fisher). Size distributions were checked on high-sensitivity DNA chips (Agilent Bioanalyzer). Samples were used to construct Nextera XT libraries (Illumina) from 100 pg of preamplified cDNA. Libraries were purified and size selected (0.5x-0.7x) using Ampure XP beads. Libraries were quantified using KAPA qPCR quantification kit (KAPA Biosystems), pooled, and sequenced using an Illumina HiSEQ 4000 instrument. Reads were mapped to the Mus musculus genome (EMSEMBL GRCm38.p4 Release 81) and ERCC sequences using GSNAP (version 2015-09-29) with parameters: -A sam –B 5 –t 24 –n 1 –Q –N 1. HTseq-count (*58*) was used to count reads mapped to each gene, with parameters: –s no. All cells with < 100,000 reads mapping to endogenous RNA and >20% reads mapping to mitochondrial genes were considered low quality and removed from downstream analyses. Data were normalized and highly variable genes were identified as previously described (*59*), using a false discovery rate threshold equal to 0.1 for the chi-squared test. Only highly variable genes were considered to perform PCA analysis, using the prcomp function in R (version 3.6.3).

UMAP was calculated using umap function (version 0.2.7) in R based on normalized expression value with 15 nearest neighbors. Differential expression analysis of old and young cells within each group was performed using DESeq2 (*60*) version 1.26.0. Pathway analysis was conducted using GSEA software v4.1.0. Gene symbols were mapped to MSigDB.v7.2.chip and overlaps with Hallmark (h.all.v7.2.symbols.gmt) gene sets were determined using the classic scoring scheme.

Data and Software Availability.

Data sets that support the findings of Dellorusso et al. have been deposited in the Gene Expression Omnibus under accession number GSE229137.

### Methods References


### Supplementary Text

Lack of PGC-1α function in mediating HSC metabolic adaptation with age

Here, we reported that *Ppargc1a*, encodes the master regulator of mitochondrial biogenesis and energy metabolism PGC-1α (*44*), was one of the most differentially expressed genes between $AT^{hi}$ and $AT^{lo}$ oHSCs (**fig. S7A**), Interestingly, *Ppargc1a* was specifically expressed in HSCs compared to the rest of the blood system, with $AT^{lo}$ oHSCs having significantly reduced *Ppargc1a* levels compared to $AT^{hi}$ oHSCs and yHSCs (**fig. S7B,C**). Moreover, transcription of *Ppargc1a* mRNA was induced following cytokine deprivation and autophagy engagement *in vitro* and repressed following cytokine-mediated proliferation in culture or genetic ablation of autophagy in $Atg12^{cKO}$ yHSCs (**fig. S7B**). To directly test the role of PGC-1α as a mediator of metabolic adaptations under autophagy-stimulating conditions, we generated $Ppargc1a^{fl/fl}$*:Mx1-Cre* ($Ppargc1a^{cKO}$) mice (*45*) to conditionally inactivate PGC-1α in HSCs, and crossed $Ppargc1a^{cKO}$ mice with *Gfp-Lc3* reporter mice to monitor autophagy levels (**fig. S7D**). In culture, $Ppargc1a^{cKO}$ HSCs showed delayed induction of autophagy 3 hours post-cytokine deprivation, which was rescued by 6 hours suggesting redundant transcriptional mechanisms to promote autophagy absent PGC-1$\alpha$. Similarly, *in vivo*, *Ppargc1a*$^{cKO}$ yHSCs, like oHSCs of either genotype, showed decreased mitotracker green (MTG) and tetramethylrhodamine-ethyl-ester (TMRE) levels compared to Ctrl yHSCs (**fig. S7E**). However, *Ppargc1a*$^{cKO}$ mice exhibited no changes in HSC numbers over time, were fully competent in regenerating the blood system following serial 5-FU-mediated myeloablation and showed normal HSC functionality upon transplantation with similarly reduced engraftment activity from both Ctrl and *Ppargc1a*$^{cKO}$ oHSCs (**fig. S7F-H**). These results demonstrate a high degree of functional redundancy in the mechanisms regulating metabolic compensation in HSC aging, with a limited role for PGC-1$\alpha$.

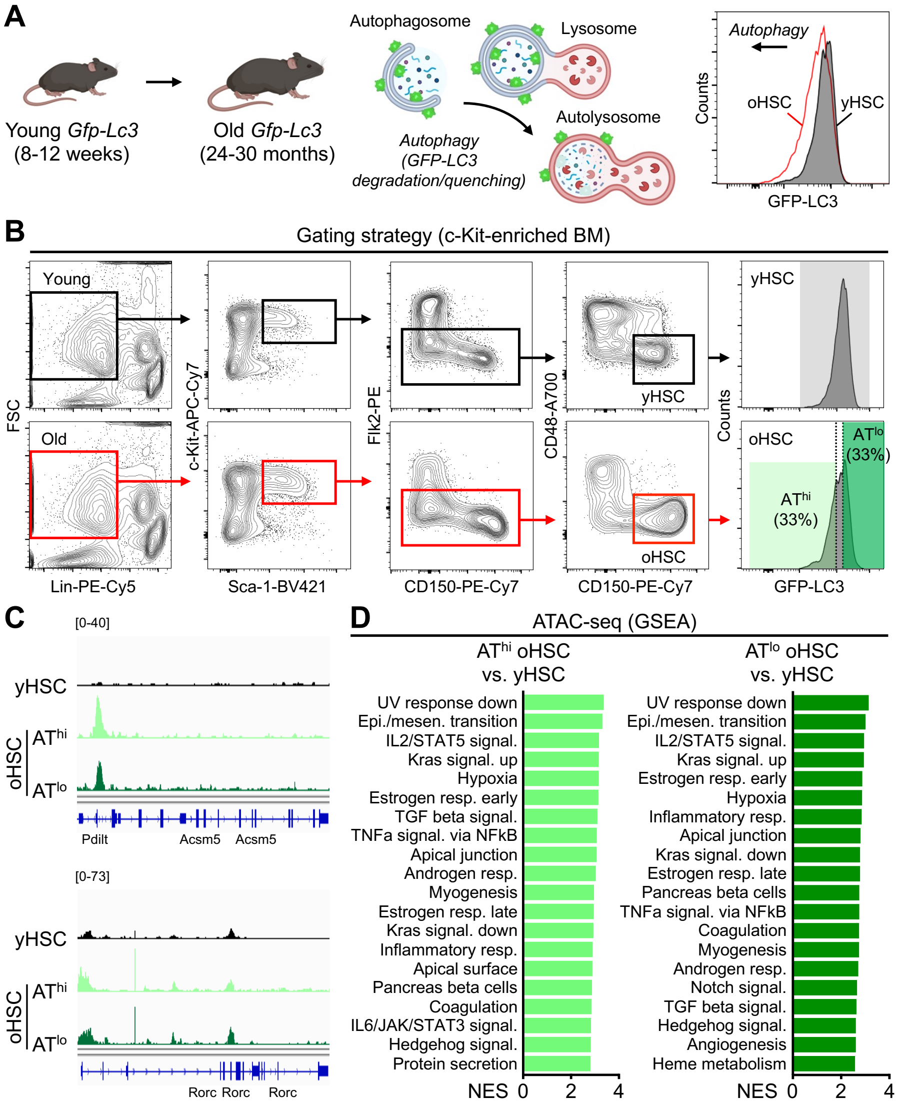


**Fig. S1. Isolation strategy and characterization of chromatin accessibility landscape.** (**A**) Schematic of GFP-LC3 reporter quenching upon lysosomal acidification and degradation and representative flow cytometry plot showing differences in reporter intensity between yHSCs and oHSCs. (**B**) Flow cytometry gating strategy for HSC isolation in young and old mice with GFP-LC3 sub-fractionation for AT$^{hi}$ and AT$^{lo}$ oHSCs. (**C**) Examples of differences in promoter proximal peak accessibility between yHSCs and AT$^{hi}$/AT$^{lo}$ oHSCs. (**D**) GSEA of the top twenty significantly (FDR q value < 0.05) differentially accessible peaks in AT$^{hi}$ oHSCs vs. yHSC (left) and AT$^{lo}$ oHSCs vs. yHSCs (right).

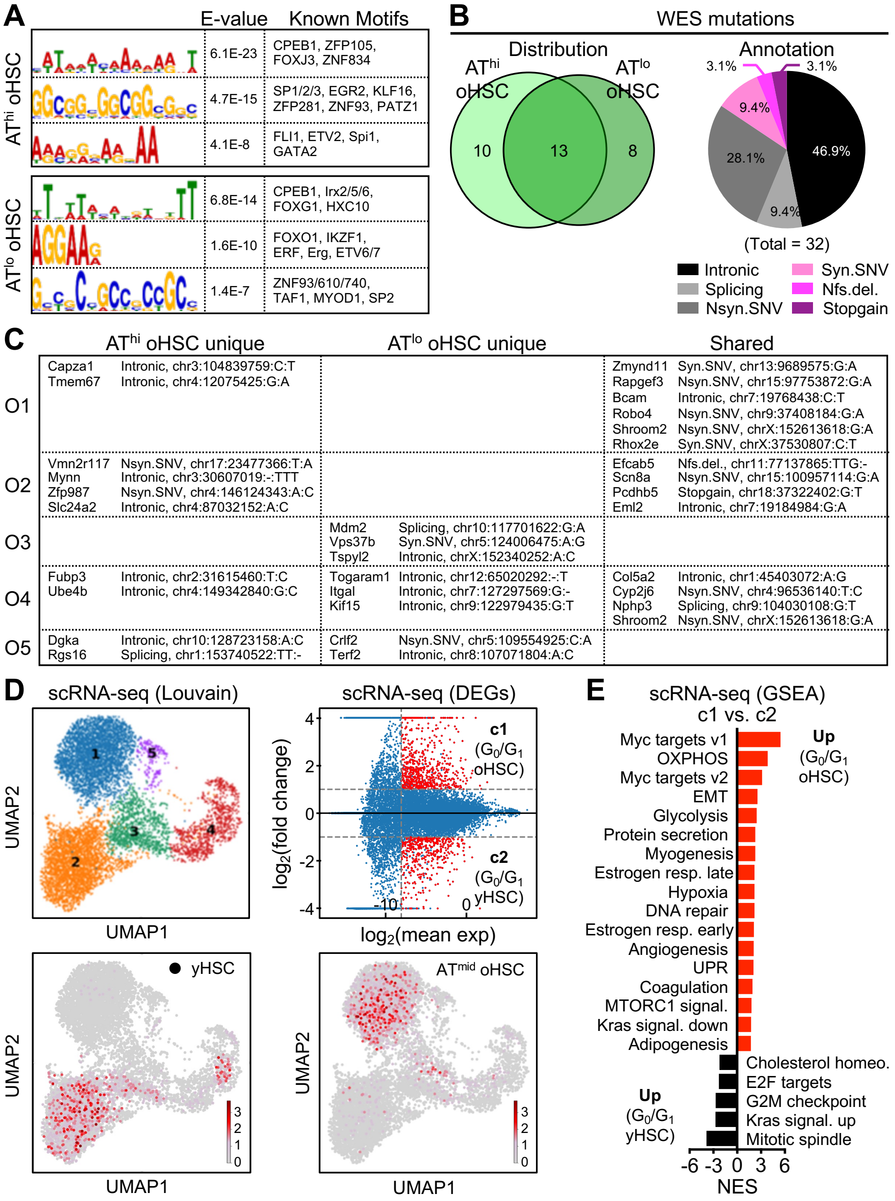


**Fig. S2. Molecular profiling of yHSCs, AT$^{hi}$ oHSCs, and AT$^{lo}$ oHSCs.** (**A**) HOMER motif analysis from the ATAC-seq dataset. (**B**) Distribution of shared and unique Mutect2 called mutations from whole exome sequencing (WES) of five biological replicates of AT$^{hi}$ and AT$^{lo}$ oHSCs, and annotated distribution of called mutations. Nsyn.SNV, nonsynonymous single nucleotide variant; Syn.SNV, synonymous single nucleotide variant; Nfs.del, non-frameshift deletion. (**C**) Table of mutations called per biological replicate. (**D**) Molecular characterization with UMAP visualization of droplet-based scRNA-seq Louvain clusters (top left), differentially expressed genes in cluster 1 (c1: $G_0/G_1$ oHSC) vs. cluster 2 (c2: $G_0/G_1$ yHSC) (top right), and projection of SMART-Seq2 transcriptomes from yHSCs (bottom left) and AT$^{mid}$ oHSCs (bottom right) onto the droplet-based scRNA-seq UMAP. (**E**) GSEA results of differentially expressed pathways between cluster 1 (c1: $G_0/G_1$ oHSC) and cluster 2 (c2: $G_0/G_1$ yHSC).

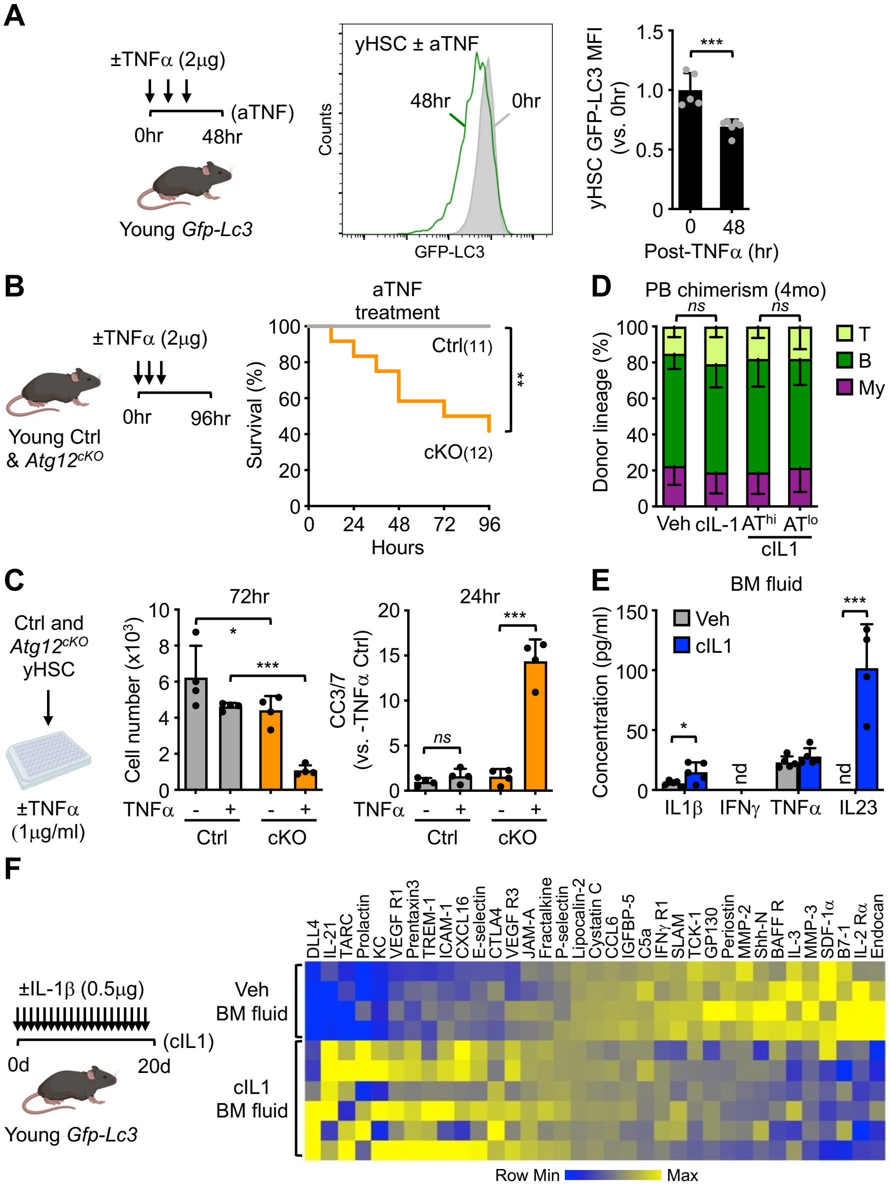


**Fig. S3. HSC autophagy response to additional inflammatory stimuli.** (**A**) Experimental design for repeated *in vivo* TNFα injections in *Gfp-Lc3* mice (left) with representative GFP-LC3 signal intensity (middle) and quantification of GFP-LC3 mean intensity fluorescence (MFI) at 0 and 48 hours (right); hr, hour. (**B**) Survival of Ctrl and *Atg12*$^{cKO}$ mice following repeated *in vivo* TNFα injections. (**C**) Survival of Ctrl and *Atg12*$^{cKO}$ HSCs upon *in vitro* TNFα exposure showing cell number (left) and cleaved caspase 3/7 (CC7/7) levels (right). (**D**) Lineage distribution in the peripheral blood (PB) at 4 months (mo) post-transplantation of the indicated HSC populations (see Fig. 2F); T; T cells; B, B cells; My, myeloid cells. (**E-F**) Differentially secreted cytokines in the BM fluid of vehicle (Veh) and chronic IL-1 (cIL1)-treated mice: (E) treatment scheme and mouse 200-Plex cytokine array measurement; and (F) mouse 20-Plex bead array measurement; nd, not detected. Data are means ± S.D.; *p ≤ 0.05, **p ≤ 0.01, ***p ≤ 0.001; *ns*, not significant.

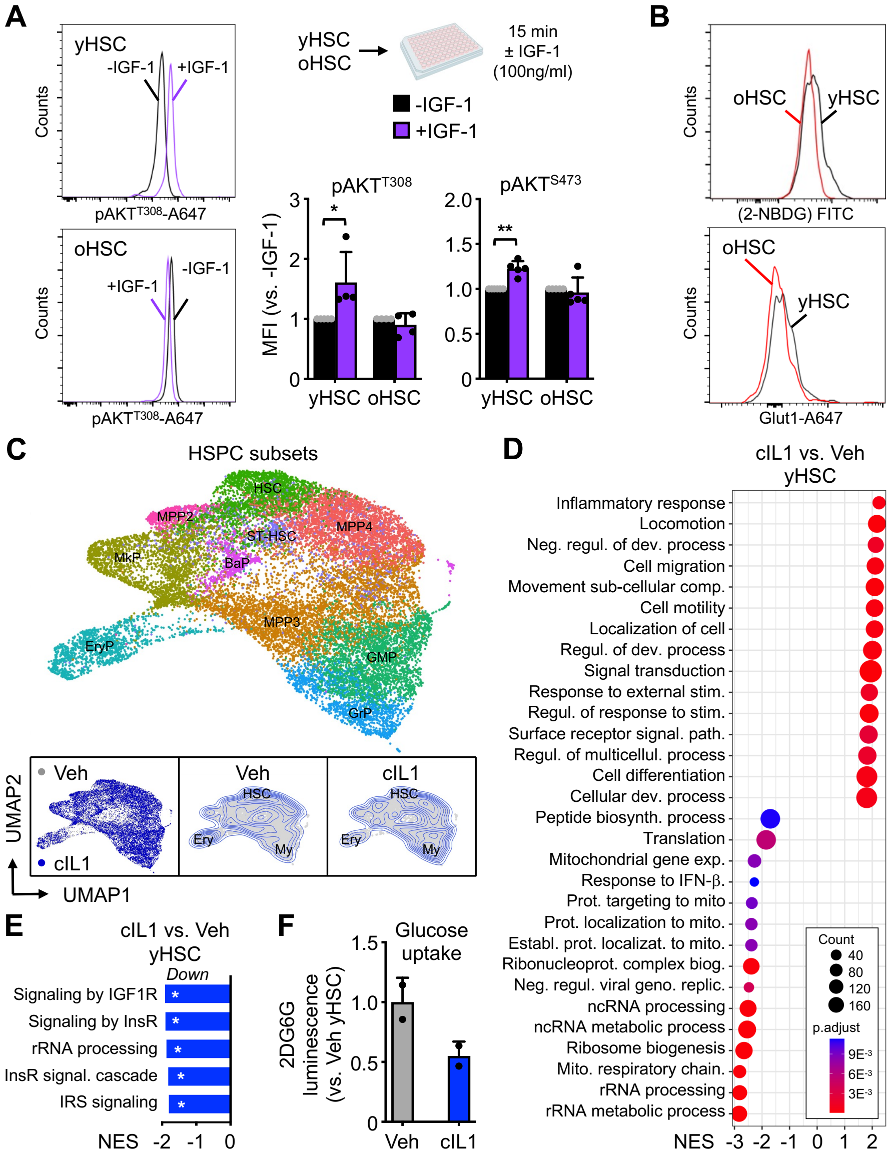


**Fig. S4. Aging and chronic inflammation impact AKT pathway activity and glycolytic metabolism in HSCs.** (**A**) Representative FACS plots (left) and pAKT$^{T308}$ and pAKT$^{S473}$ MFI quantification (right) in yHSCs and oHSCs following acute 15 minutes (min) stimulation with or without (±) 100 ng/ml IGF-1 in liquid culture. (**B**) Representative FACS plots for 2-NBDG and Glut1 MFI quantification in young and old HSCs. (**C**) UMAP visualization of droplet-based scRNA-seq analyses of mixed LK/LSK BM fractions isolated from vehicle (Veh, 4 pooled mice) or cIL1 (6 pooled mice) treated mice. Results show Louvain clusters on integrated UMAP (top), and contour plots split by group, Veh (9,009 cells) and cIL1 (11,168 cells) (bottom left). (**D**) GSEA of top 15 significantly enriched and suppressed pathways by normalized enrichment score (NES) in cIL1 vs. Veh HSC scRNA-seq dataset. (**E**) Reactome pathway analysis of bulk RNA-seq from cIL1 vs. Veh HSC RNA-seq dataset (GSE165810) (*33*). (**F**) Glucose uptake in Veh and cIL1 HSCs as measured by 2-deoxy-D-glucose-6-phosphate (2DG6P) luminescence after 4 hours of culture. Data are means ± S.D. except when indicated (E, ± S.E.M.); $*p \leq 0.05$, $**p \leq 0.01$.

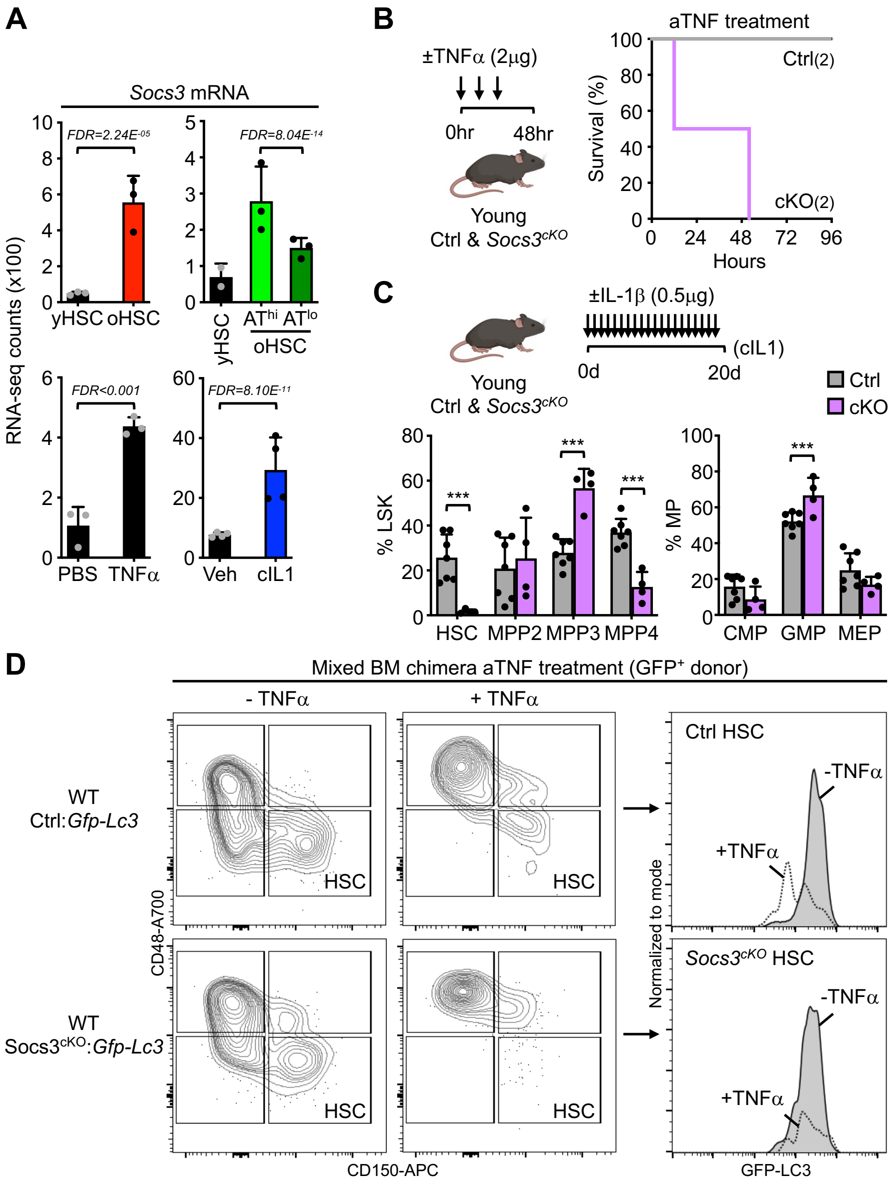


**Fig. S5. Role of Socs3 in mediating HSC inflammatory response.** (**A**) *Socs3* mRNA transcript levels from the indicated RNA-seq datasets; yHSC vs. oHSC (top left), yHSC vs. AT$^{hi}$ oHSC vs AT$^{lo}$ oHSC (top right), PBS vs. TNFα yHSC (bottom left), Veh vs. cIL1 yHSC (bottom right; GSE165810) (*33*); FDR, false discovery rate. (**B**) Survival of Ctrl and *Socs3$^{cKO}$* mice following acute TNFα (aTNF) treatment. (**C**) Immunophenotyping of Ctrl and *Socs3$^{cKO}$* mice following chronic IL-1β treatment with HSCs and multipotent progenitors (MPP) 2 to 4 expressed as percent of LSK (Lin$^-$/c-Kit$^+$/Sca-1$^+$) BM cells and myeloid progenitors as percent of MP (Lin$^-$/c-Kit$^+$/Sca-1$^-$) BM cells. CMP, common myeloid progenitor; GMP, granulocyte/macrophage progenitor; MEP, megakaryocyte/erythrocyte progenitor. (**D**) Representative FACS plots showing the gating of GFP$^+$ donor HSCs in Ctrl and *Socs3$^{cKO}$* mixed BM chimera, and changes in GFP-LC3 signal intensity following TNFα exposure. Data are means ± S.D. except when indicated (**A**, FDR); ***p ≤ 0.001.

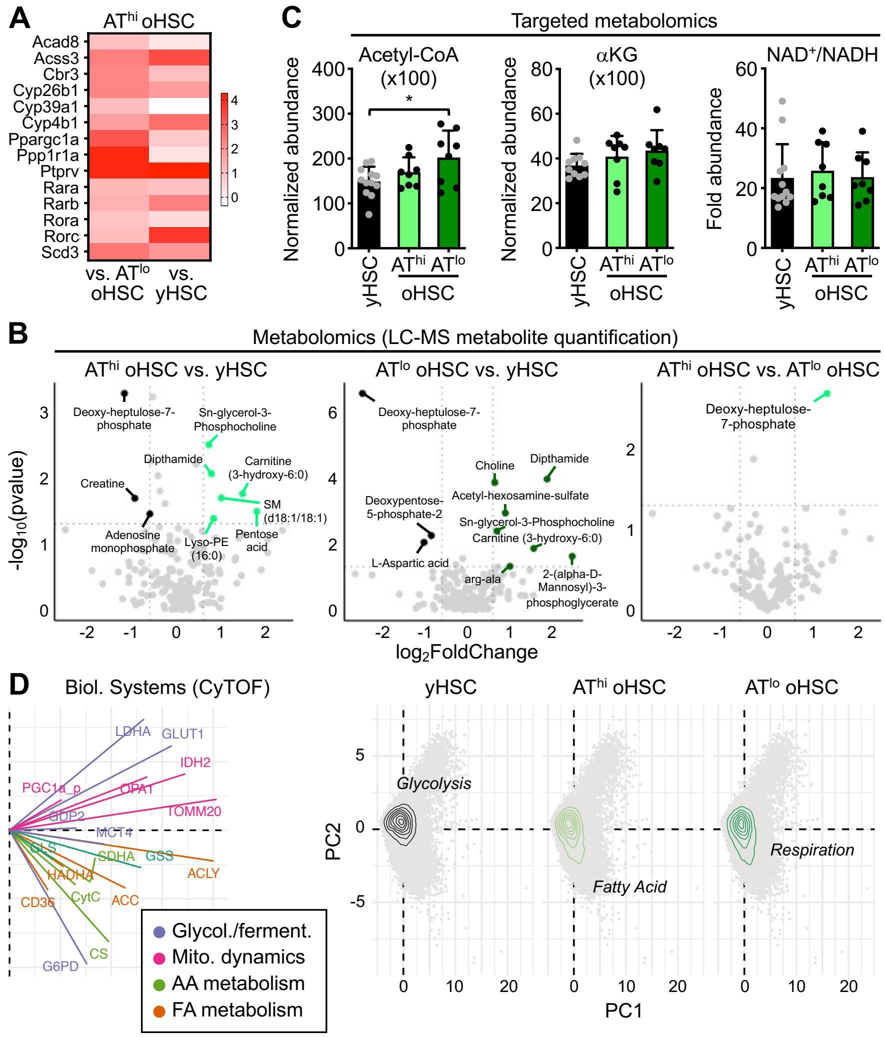


**Fig. S6. Metabolic profiling of yHSCs, AT$^{hi}$ oHSCs, and AT$^{lo}$ oHSCs.** (**A**) Heatmap of differentially expressed genes in AT$^{hi}$ oHSC vs. AT$^{lo}$ oHSC or yHSC bulk RNA-seq showing fold change in selected metabolic enzymes and transcriptional regulators involved in nutrient metabolism. (**B**) Differentially abundant metabolites for the indicated pairwise comparisons following low-input shotgun metabolomics. Results are shown with an exploratory threshold of logarithmic fold change (LFC) 0.6 and p <0.10; LC-MS, Liquid chromatography-mass spectrometry. (**C**) Levels of acetyl-CoA, alpha-ketoglutarate (αKG), and NAD$^{+}$/NADH ratio in the indicated HSC populations following targeted metabolomics profiling. Data are means ± S.D.; *p ≤ 0.05. (**D**) Principal components (PC) loadings of mass cytometry (CyTOF) metabolic regulome panel and expression density highlighting the major metabolic pathway over-represented in yHSC, AT$^{hi}$ oHSC, and AT$^{lo}$ oHSC. Metabolic pathway classification of protein targets in different biological (Biol.) systems is indicated. Glycol., glycolysis; Ferment., fermentation; Mito., mitochondria; AA, amino acid; FA, fatty acid.

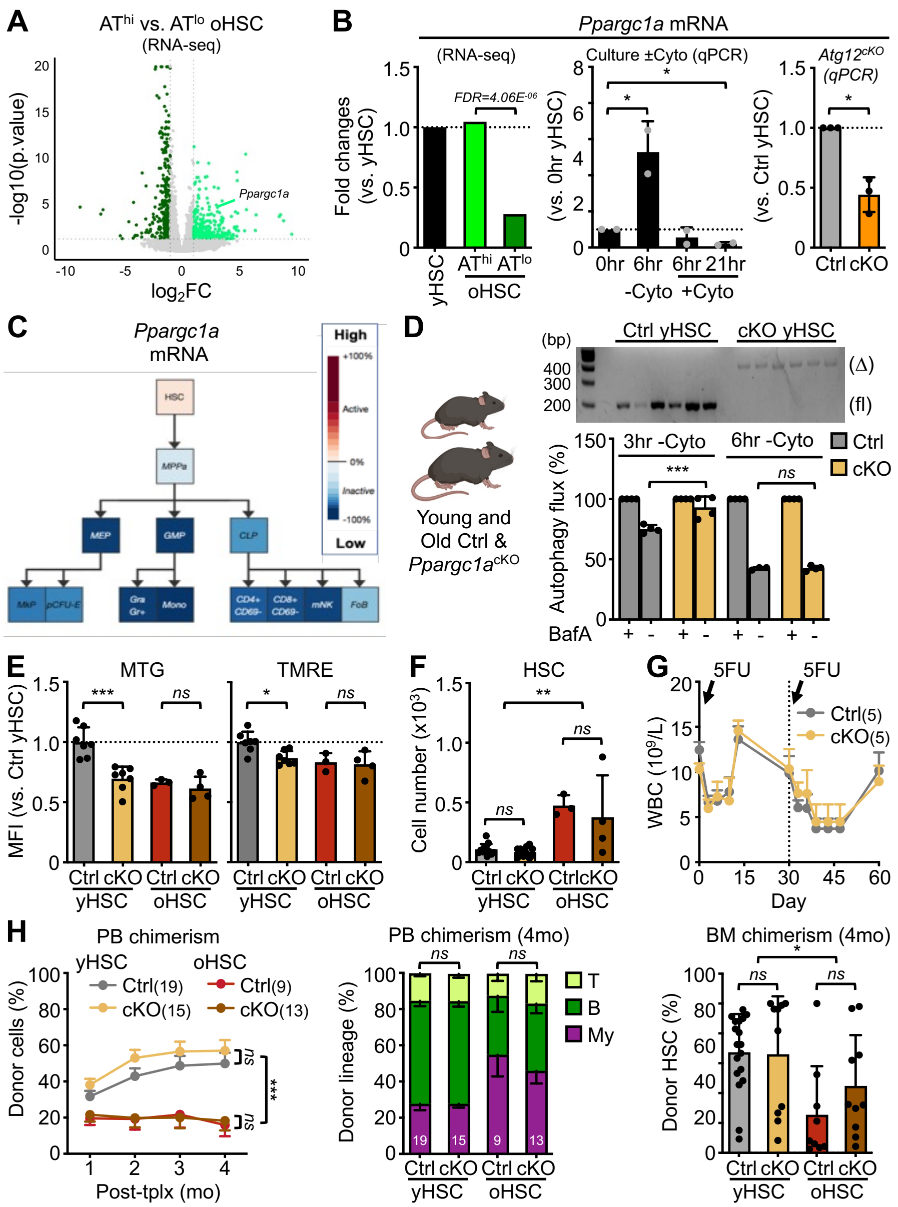


**Fig. S7. Lack of PGC-1α function in mediating HSC metabolic adaptation with age.** (**A-B**) *Ppargc1a* mRNA expression in the indicated RNA-seq and qRT-PCR (qPCR) datasets: (A) Volcano plot visualization; and (B) relative fold changes; hr, hour; ±Cyto, culture with or without cytokines; Ctrl, control; cKO, *Atg12*$^{cKO}$. (**C**) *Ppargc1a* expression in the mouse hematopoietic hierarchy (GSE 34723). (**D**) Experimental scheme for deleting *Ppargc1a* in adult mice (left) with deletion efficiency in single HSC colony PCR (top) and autophagy flux following cytokine deprivation ± Bafilomycin A (BafA, bottom). (**E-H**) Characterization of Ctrl and *Ppargc1a* conditional knockout (*Ppargc1a*$^{cKO}$ or cKO) young and old mice: (E) MTG and TMRE HSC levels; (F) HSC cellularity; (G) blood regeneration following serial 5-FU injection (WBC, white blood cell counts); and (H) regenerative capacity of the indicated HSC populations following transplantation (tplx) into lethally irradiated recipients (250 HSC/recipient). Results show overall engraftment in peripheral blood (PB) over time (left) with lineage distribution (middle) and HSC chimerism (right) at 4 months (mo) post-tplx. Data are means ± S.D. except when indicated (G and H, ± S.E.M.); *p ≤ 0.05, **p ≤ 0.01, ***p ≤ 0.001; *ns*, not significant.

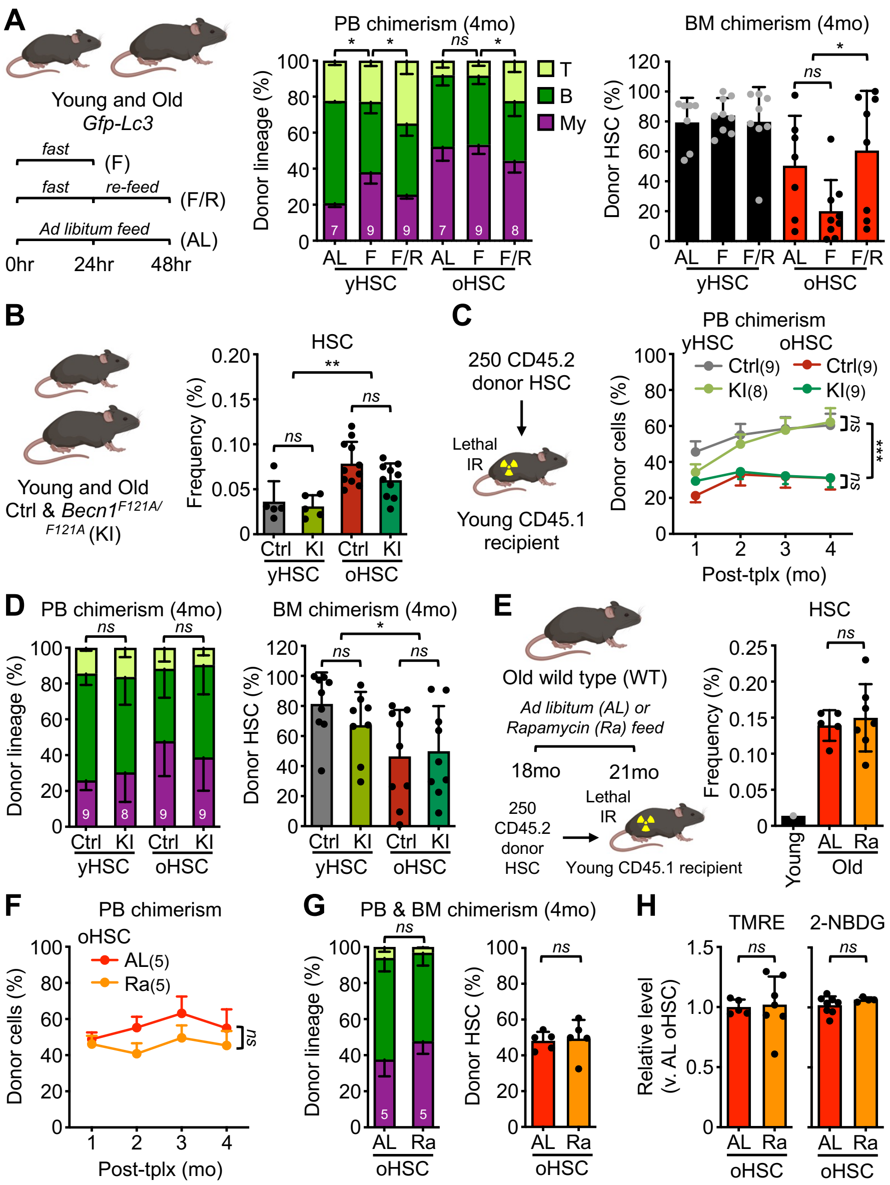


**Fig. S8. Impact of chronic autophagy engagement on oHSC fitness.** (**A**) Engraftment results for the indicated HSCs (left) showing lineage distribution (middle) and HSC chimerism (right) at 4 months (mo) post-transplantation (tplx). Results are related to experiment described in Figure 4C. (**B-D**) Constitutive autophagy engagement in young and old *Becn1* knockin (KI) vs. control (Ctrl) mice: (B) HSC BM frequencies; (C) overall engraftment in peripheral blood (PB) over time, and (D) lineage distribution (left) and HSC chimerism (right) at 4 months post-transplantation of the indicated HSC populations into lethally irradiated recipients. (**E-H**) Constitutive autophagy activation in rapamycin (Ra) feed vs. Ad Libitum (AL) feed old mice: (E) experimental scheme (left) and HSC BM frequencies (right) compared to young mice; (F) overall engraftment in PB over time and (G) lineage distribution (left) and HSC chimerism (right) at 4 months post-transplantation of the indicated HSC populations into lethally irradiated recipients; and (H) relative TMRE and 4 hours ex-vivo 2-NBDG uptake in the indicated HSC populations. Data are means ± S.D. except when indicated (C and F, S.E.M.); *p ≤ 0.05, **p ≤ 0.01, ***p ≤ 0.001; *ns*, not significant.

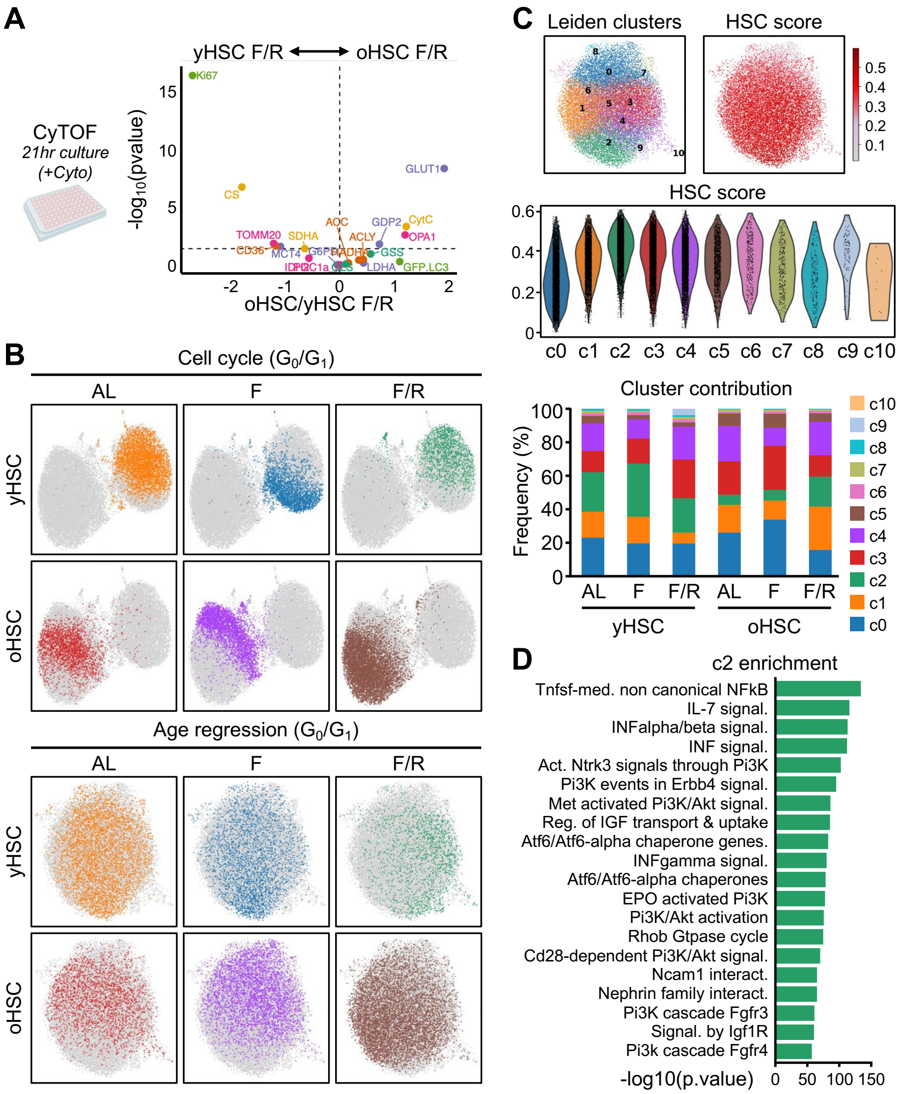


**Fig. S9. Single-cell RNA-seq profiling of young and old HSCs following fasting and refeeding.** (**A**) Differentially expressed metabolic proteins in F/R yHSC vs. F/R oHSC following 21hr culture in full cytokine media. (**B**) UMAP visualization of all predicted $G_0/G_1$ cells with cell labeling by sample type (top) and upon regression for age and feeding pattern (bottom); AL, Ad Libitum; F, fast; F/R, fast/refeed. Results are related to experiment described in Figure 4F. (**C**) Leiden clusters observed at a resolution of 1 following age and feeding pattern regression (top left) and cell HSC score (top right) with violin plot of HSC score by cluster (middle) and Leiden cluster representation by sample type (bottom). (**D**) GSEA results for Reactome pathway analyses of cluster 2 (c2 vs. all). The full list of GSEA results per cluster is presented in table S2.

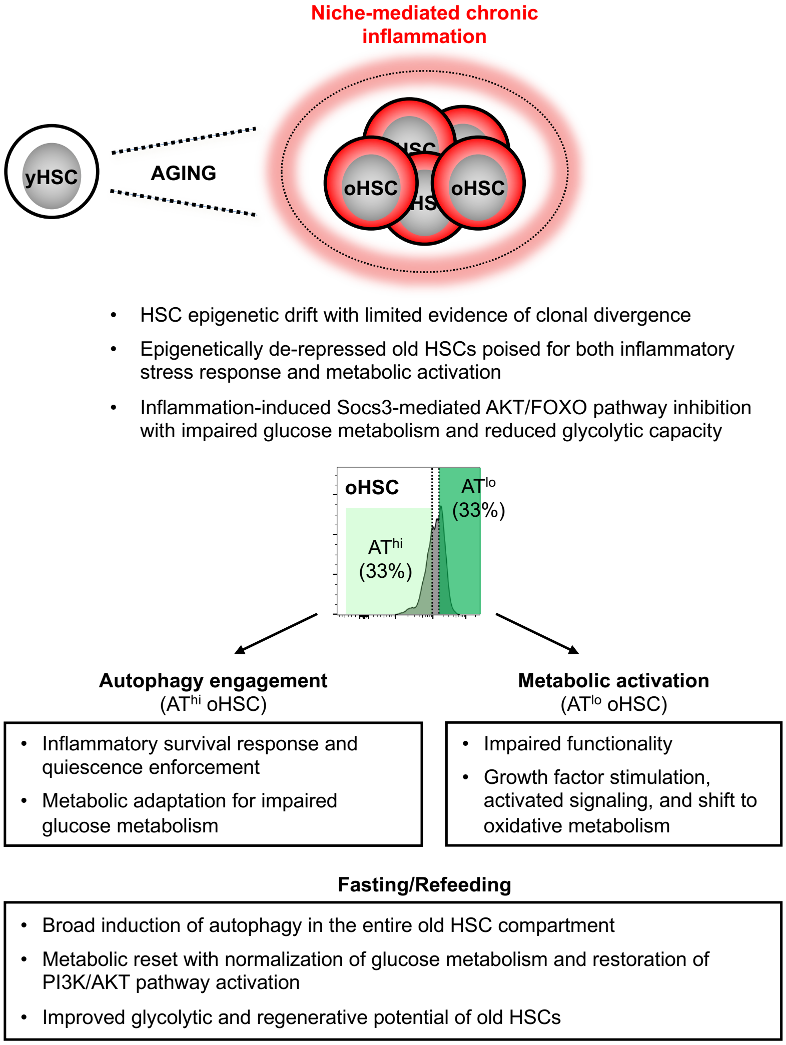


**Fig. S10. Model for autophagy in old HSCs as an adaptive response to chronic niche inflammation, and a target for metabolic reset and functional restoration.** With age, the entire HSC compartment displays epigenic drift with more accessible chromatin regions for genes involved in inflammatory stress response and altered metabolism. However, we found limited evidence of clonal divergence at the epigenetic or genetic levels in old HSCs. Instead, chronic inflammation from the aging BM niche microenvironment led to a broad suppression of PI3K/AKT signaling network via Socs3 upregulation, resulting in suppressed glycolytic metabolism in the entire HSC compartment and with compensatory activation of autophagy in old HSCs with the highest inflammatory response. In this context, autophagy is a cytoprotective and quiescence-enforcing response to inflammation allowing metabolic adaptation for impaired glucose metabolism. In its absence, old HSCs shift to oxidative metabolism, enter the cell cycle, and lose their regenerative capacity. Strikingly, acute fasting/refeeding intervention induces a broad activation of autophagy and improvement of glycolytic metabolism in the entire old HSC compartment, leading to functional restoration and increased regenerative potential. These results show that modulating the autophagy axis enables a metabolic reset of old HSCs for demand-adapted fitness.

**Table S1. Mass Cytometry Antibody Staining Panel.** Metal-isotope labeled antibodies utilized for HSPC isolation and metabolic protein expression are listed. For each antibody target, corresponding clones and providers are listed, along with the conjugated isotope and mass channel it was detected in. Staining concentrations for each antibody in the panel are also provided.
*(Separate document)*

**Table S2. Pathway analysis of young and old HSCs single-cell RNA-seq clusters following fasting/refeeding.** Leiden clusters identified in $G_0/G_1$ young and old HSCs following age and feeding pattern linear regression (shown in Fig S9C) were analyzed by GSEA for unique gene expression pathway enrichment in each cluster compared to all others. For each cluster, the top twenty enriched Reactome pathways are listed with corresponding enrichment scores, p values, and FDR adjusted p values.
*(Separate document)*
